# Epigenetic Disordering Drives Stemness, Senescence Escape and Tumor Heterogeneity

**DOI:** 10.1101/2024.12.29.629346

**Authors:** Elena Magnani, Filippo Macchi, Tijana Randic, Charlene Chen, Bhavani Madakashira, Shashi Ranjan, Sema Elif Eski, Sumeet P. Singh, Kirsten C. Sadler

## Abstract

Tumor heterogeneity is the substrate for tumor evolution and the linchpin of treatment resistance. Cancer cell heterogeneity is largely attributed to distinct genetic changes within each cell population. However, the widespread epigenome repatterning that characterizes most cancers is also highly heterogenous within tumors and could generate cells with diverse identities and malignant features. We show that high levels of the epigenetic regulator and oncogene, UHRF1, in zebrafish hepatocytes rapidly induced methylome disordering, loss of heterochromatin, and DNA damage, resulting in cell cycle arrest, senescence, and acquisition of stemness. Reducing UHRF1 expression transitions these cells from senescent to proliferation-competent. The expansion of these damaged cells results in hepatocellular carcinomas (HCC) that have immature cancer cells intermingled with fibroblasts, immune and senescent cells expressing high UHRF1 levels, which serve as reservoirs for new cancer cells. This defines a distinct and heterogenous HCC subtype resulting from epigenetic changes, stemness and senescence escape.

## Introduction

Tumor suppression is achieved through the coordination of cell death, senescence and immune clearance of pre-cancerous cells. Cancer forms through the selective advantage of those cells that successfully overcome these mechanisms and acquire the ability to garner resources to proliferate and to survive stress imposed by the microenvironment. In most tumors, these features are not all encompassed by a single cancer cell, but the tumor as a whole evolves as an ecosystem made up of cells with a range of identities and capacities that sustain tumor survival and growth^1^. This heterogenous nature of tumors presents a major challenge in developing effective therapies. Indeed, in many cases, relapse is caused by treatment resistance in small population of cancer cells^2^.

DNA damage, mutations, and genomic rearrangements create a foundation for tumor evolution and progression. While cells with excessive DNA damage or aneuploidy that substantially impair cellular function are typically eliminated through cell death, others with less damage such as induced by a very high level of oncogene expression undergo a type of senescence that both restricts their expansion and sends signals for immune mediated clearance^3–5^. Importantly, damaged cells that fail to be cleared by the immune system^6^ or which have a lower level of oncogene overexpression^3^ evade tumor suppression, acquire mechanisms to either outcompete their neighbors for resources or kill them off^7^, and to survive in conditions that are suboptimal for non-transformed cells^1,3,8,9^.

While mutational diversity is a major mechanism that drives cancer cell selection, the widespread epigenetic alterations found in nearly all cancers offer an alternative route to reshaping gene expression and cellular identity to create heterogenous tumors. The cancer epigenome is characterized by a disordered pattern of DNA methylation and changed chromatin accessibility. The DNA methylation pattern of healthy cells is characterized by heavily methylated repeats, transposable elements, and gene bodies while promoters are unmethylated. In cancer, this pattern becomes randomized, with CpGs in some promoters gaining methylation and those in other genomic elements losing methylation in a random pattern^10,11^. These changes occur early in tumorigenesis^12,13^ and are induced by cancer causing insults such as aging^14^ or viral infection^15^. In some cases, DNA methylation and chromatin accessibility changes can directly regulate genes that contribute to cancer. For instance, in prostate cancer, DNA methylation changes exert long range effects across the genome to activate oncogenes and other cancer drivers^16^ and in melanoma, widespread changes to the chromatin landscape are required for responding to subsequent oncogenic stimuli^17^. In other cases, DNA methylation loss drives mutations and chromosomal instability^18–22^, shaping the mutational landscape that drives tumor evolution.

Whether epigenetic changes alone are sufficient to cause cancer, or if they are instead a mechanism that causes chromosomal instability as a mechanism of transformation is important to resolve, since, unlike genomic rearrangements, epigenetic changes hold the possibility of manipulation and reversion to normal states. Recent studies show that epigenetic changes on their own^23^ or in combination with environmental stimuli^24^ can cause cancer without mutations or other genomic changes. Our work demonstrating that overexpression of the epigenetic regulator ubiquitin, PHD and ring finger domain containing protein 1 (UHRF1) in zebrafish hepatocytes causes hepatocellular carcinoma (HCC)^25^ provided functional validation that an epigenetic regulator can be an oncogene. UHRF1 along with DNA methyltransferase 1 and other DNA modifying enzymes regulate maintenance DNA methylation^26–29^, recruits histone methyltransferases that deposit repressive histone modifications^30,31^, and is required for repair of double strand breaks^31,32^. This further supports the hypothesis that an epigenetic regulator of DNA methylation can be a cancer driver. Whether the epigenetic damage caused by UHRF1 overexpression causes DNA damage is important to examine since the promise of reversing epigenetic changes to treat cancer will be most effective if the changes do not induce the irreversible genomic changes that characterize many treatment-refractory cancers.

It is well established that precancerous lesions and senescent cells have epigenetic changes that are similar to cancer^12^. Senescence was previously considered a homogenous cellular state caused by replicative exhaustion, aging or other extreme forms of stress that renders cells either dysfunctional or beyond repair. It is now understood to be a complex phenotype that has different features depending on the cause, cellular context or duration. DNA damage is the most common trigger, and thus a persistent DNA damage response (DDR) and the resulting activation of *tp53* is a common feature of senescent cells^33,34^. The epigenetic and chromatin changes are caused by some senescent stimuli include formation of senescence associated heterochromatin foci (SAHF), and dramatic changes to the distribution of DNA methylation and other heterochromatin marks^33,35^. Senescent cells have a proinflammatory senescent associated secretory phenotype (SASP) that invokes an immune response that can clear senescent cells and, in many cases induce senescence in neighboring cells^36^. If immune clearance is incomplete, persistent senescent cells can contribute to chronic inflammation that can cause tissue damage and be tumorigenic^37^.

Several studies have demonstrated that the epigenetic landscape in precancerous lesions and in senescent cells have similarities to cancer^12,13^. In addition, stemness is a feature of some senescent cell states^38,39^, which may enable their escape. Together, these features prime senescent cells to be the precursor for malignancy. Moreover, given that epigenetics is a main mechanism by which cells acquire and sustain identity, it is feasible that the epigenetic damage which induces senescence can also underlie the heterogenous tumors once these cells escape.

To identify the mechanism of epigenetic change mediated senescence induction, escape and tumor heterogeneity, we utilize zebrafish with high UHRF1 overexpression in hepatocytes (*Tg(fabp10a:hUHRF1-EGFP)high*, hereafter referred to as *hUHRF1-high*). This model enables real-time analysis of epigenetically induced senescence within 2 days and cancer formation within 2 weeks of UHRF1 overexpression, offering a unique system to identify the mechanisms underlying epigenetic changes leading to senescence, escape, and hepatocarcinogenesis. We hypothesize that epigenetic dysregulation driven by UHRF1, either alone or in combination with DNA damage, induces a senescent and stem-like state, with some cells evading clearance to undergo malignant transformation and drive tumor heterogeneity. We identify methylome disordering, transposon activation, DNA damage and Atm-tp53 activation as the mechanism of hUHRF1 driven senescence and find that this is accompanied by the acquisition of stemness which enables senescence escape, expansion of cells with epigenetic damage and generation of highly heterogeneous HCC.

## Results

### UHRF1 overexpression causes double strand breaks and cellular senescence

Senescence is a pleiotropic cell state that has a range of definitive features including, cell cycle withdrawal, β-galactosidase (SA-β-gal) staining, DNA damage and a persistent DDR characterized by tp53 and p21 activation, SASP, SAHF and other cellular and molecular markers^40^. The combination of these features varies depending on the type of senescence^5,41^. We previously showed that hUHRF1 overexpression in zebrafish hepatocytes is turned on by 72 hpf, during hepatocyte differentiation, and results in reduced proliferation at 120 hpf, strong SA-β-gal staining in the liver of nearly all larvae at 120 hpf, but not at 96 hpf and induced *tp53* signaling^25^. Since senescence is pleotropic, we established the features of UHRF1-induced senescence and the time course of these features by assessing cell proliferation, *tp53* activation based on gene expression, DNA damage and the DDR, SASP and immune cell recruitment and SAHF (Figure 1A). The zebrafish liver grows rapidly between 72-120 hpf, as demonstrated by a high percent of cells undergoing DNA replication as marked by EdU incorporation over 25%, 15% and 2% of cells in control livers at 80, 96 and 120 hpf respectively. This was reduced to 15% and 5% in hUHRF1- high larvae at 80 and 96 hpf, respectively, and there was no difference at 120 hpf (Figure 1B). Using nuclear morphology to identify hepatocytes (Figure 1C) we found that while both hepatocytes and non-hepatocytes incorporate EdU in control livers at 96 hpf, virtually no hepatocytes incorporated EdU in hUHRF1-high livers (Figure 1C-D). We eliminated the possibility that the transgene insertion caused senescence due to a bystander effect by showing that CRISPR-Cas9 mutagenesis of the transgene (Figure S1A-B) rescued SA-β-gal staining (Figure S1C). We mapped the insertion of the transgene to Chromosome 14 and RNAseq analysis of genes at this locus (Figure S1D) showed them to be unaffected in hUHRF1-high livers at 80 or 120 hpf (Figure S1E).

**Figure 1.**
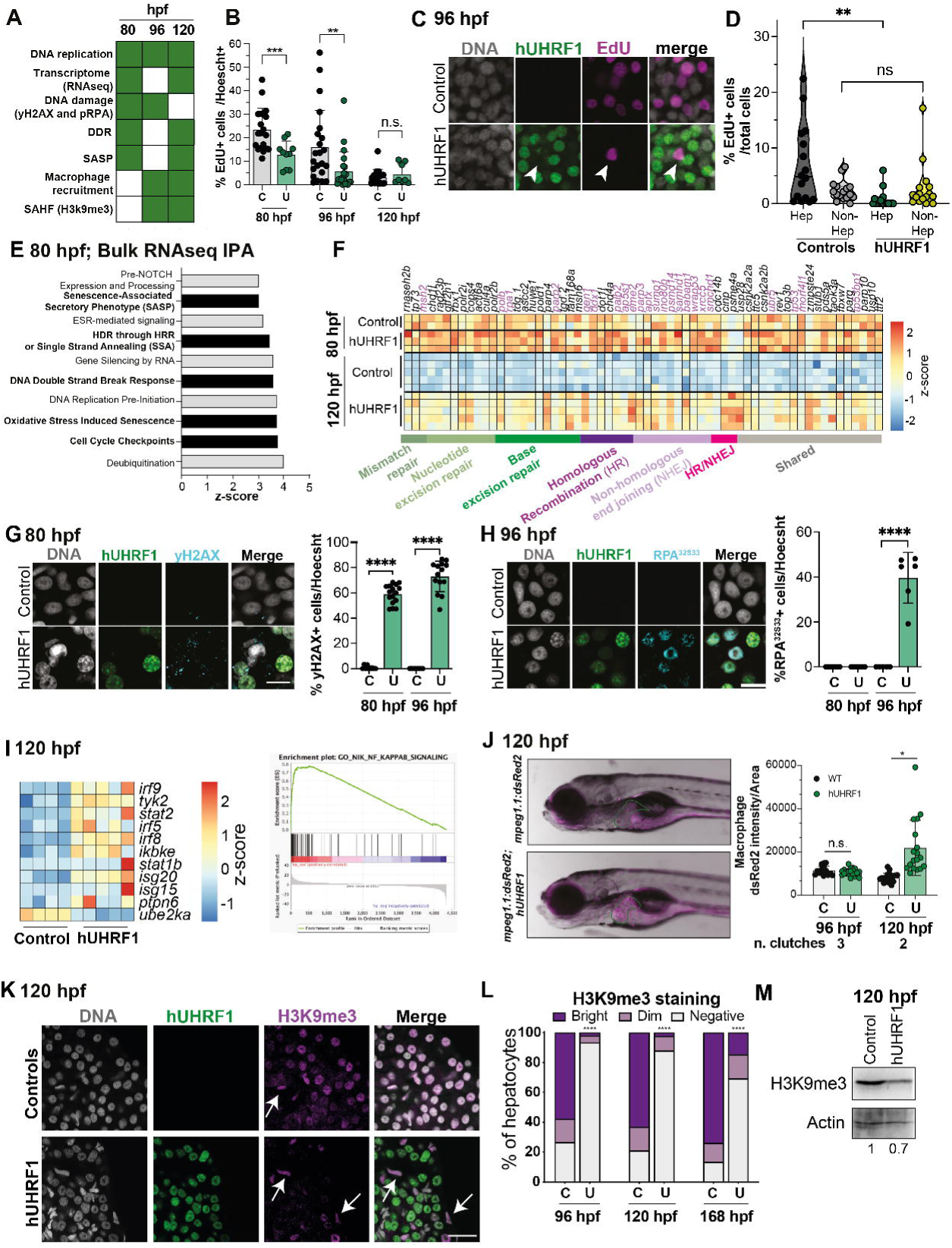
UHRF1 overexpression causes double strand breaks and cellular senescence. **A.** Scheme indicating the assays performed to characterize UHRF1-induced senescence**. B.** EdU incorporation assay of hUHRF1-EGFP and controls livers at 80, 96, and 120 hpf. Significance is measured by unpaired t-test. *** means p-values < 0.001, ** means p-values < 0.01. **C.** Representative images of EdU signal in hUHRF1 and Controls livers at 96 hpf. Arrow indicates a EdU positive hepatocyte negative for hUHRF1. **D.** Quantification of EdU positive hepatocytes and non-hepatocytes separated by nuclear morphology at 96 hpf in hUHRF1-EGFP and control livers. Significance is measured by unpaired t-test. ** means p-values < 0.01. **E.** Ingenuity Pathways Analysis of differentially expressed genes (padj < 0.05) at 80 hpf in hUHRF1 compared to Control livers. In black the pathways associated to senescence. **F.** Heatmap of DNA damage repair genes (DNA repair, GO:0006281) differentially upregulated (padj < 0.05 and log2 Fold Change > 0) at 80 hpf or 120 hpf in hUHRF1 compared to controls. Genes directly involved in double strand break repair are labelled in purple. **G.** Representative images of immunofluorescence assay of yH2AX in hUHRF1-EGFP and control at 80 hpf. Quantification shows the percentage of yH2AX positive cells at 80 and 96 hpf. Significance is measured by unpaired t-test. **** means p-values < 0.0001. **H.** Representative images of immunofluorescence assay of phosphorylated RPA (pRPA) in hUHRF1-EGFP and control livers at 96 hpf. Quantification shows the percentage of pRPA positive cells at 80 and 96 hpf. Significance is measured by unpaired t-test. ** means p-values < 0.01. **I.** Heatmap of a subset of SASP genes of hUHRF1 compared controls at 120 hpf and enrichment plot of NFkB pathway in hUHRF1 compared controls at 120 hpf. **J.** Representative images of macrophage reporter line (mpeg1.1:dsRed2) in hUHRF1 and control fish at 5 dpf. In purple the macrophages, in green the outline of the liver. Quantification of macrophage recruitment measured as average intensity normalized on liver area at 96 and 120 hpf. Significance is measured by unpaired t-test and * means p-value < 0.05. **K.** Representative images of immunofluorescence of H3K9me3 in hUHRF1 and controls at 120 hpf. **L.** Quantification of H3K9me3 staining at 96, 120 and 168 hpf. **M.** Western blot of H3K9me3 of controls and hUHRF1 livers at 120 hpf. Numbers indicates levels of H3K9me3 normalized on H3 intensity. For each experiment at least 3 biological replicates are used.

Bulk RNAseq of the liver of hUHRF1-high compared to controls at 80 hpf, the earliest timepoint where senescence features were identified, showed 51 genes significantly upregulated (padj < 0.05 & log2 Fold Change > 0) and 394 downregulated (padj < 0.05 & log2 Fold Change < 0, Supplemental Figure 2A-B, Supplemental Table S1), with the majority of the genes that were differentially expressed at 80 persisting to 120 hpf (Supplemental Figure 2C, Supplemental Table S1). Ingenuity Pathway Analysis (IPA) of DEGs found that the upregulated genes were enriched in pathways involved in SASP, cell cycle checkpoints and the response to double stranded breaks (DSB) (Figure 1E), indicating activation of the DDR. Comparing all DDR genes upregulated at 120 hpf showed that many genes, including *tp53* and several genes involved in DSB repair, were induced at 80 hpf and persisted to 120 hpf (Figure 1F). The persistence of DNA damage was shown by strong staining of ©H2AX foci which mark sites of DNA damage in hUHRF1-high hepatocytes as early as 80 hpf and persisted to 96 hpf but were absent from control hepatocytes at all time points (Figure 1G). Phosphorylated replication protein A (p-Rpa) accumulates on single stranded DNA generated during DSB repair and was first detected in hUHRF1-high samples at 96 hpf (Figure 1H), following γH2AX detection. Together, these data show that DNA damage is an early response to UHRF1 overexpression and suggests that DSB is the main form of damage caused by hUHRF1 overexpression.

We assessed whether SASP and immune cell recruitment was a feature of hUHRF1 induced senescence by analyzing bulk RNAseq data from 120 hpf (Figure 1I) and counting the number of macrophages in the liver using the transgenic marker *Tg(mpeg1:mCherry)*. Many SASP genes and a key upstream immune regulator, NFkB, were induced in hUHRF1-high livers (Figure 1I) and the number of macrophages in the liver were significantly increased at 120 hpf but not at 96 hpf (Figure 1J), indicating a classical SASP mediated immune response.

SAHF is a common feature of oncogene induced senescence^33,42^ and is marked by histone H3 trimethylated lysine 9 (H3K9me3). There were H3K9me3 puncta in the nucleus of nearly all liver cells in controls and was completely absent from hepatocytes in UHRF1-high livers but was retained in non-hepatocytes (Figure 1K) as early as 96 hpf livers and persisted to 168 hpf (Figure 1L). Western blotting confirmed the loss of this epigenetic mark in hUHRF1-high livers (Figure 1M).

These data demonstrate that hUHRF1 overexpression rapidly induces a sequence of senescence features, starting with DNA damage, DDR with Tp53 activation and SASP by 80 hpf, followed by macrophage recruitment, pRPA and then SA-β-gal within 2 days of overexpression. Surprisingly, these senescent cells are depleted of H3K9me3, suggesting that hUHRF1 induces widespread defects in the repressive epigenome which causes DNA damage.

### Heterochromatin loss and TE activation as a mechanism of hUHRF1 induced DNA damage

Senescent cells have widespread changes to the pattern of DNA methylation, including partially methylated domains (PMDs) that reflect disordering of methylation; a feature that is common between senescent and cancer cells^12^. Given the striking depletion of H3K9me3 caused by hUHRF1 overexpression and the important role of UHRF1 in methylome patterning, we assessed DNA methylation in the liver of hUHRF1-high livers. 5mC was prominent at the nuclear lamina in control hepatocytes and was found in the nucleoplasm in hUHRF1 larvae at 96 hpf, and then completely repatterned to mirror the pattern of hUHRF1-EGFP distribution in the nucleus at 120 hpf (Figure 2A). Methylome analysis using reduced representation bisulfite sequencing (RRBS) of the liver at 120 hpf shows that nearly half of all CpGs analyzed had a significant difference in methylation levels, with 11.8% of CpGs changed methylation levels more than 25% (Figure 2B). Interestingly, this change is randomized, with both gain and loss of methylation distributed across every chromosome (Figure 2C) without any clear differentially methylated domains, suggestive of PMDs (Figure 2D). By analyzing only those CpGs that had a change in methylation of >25% in hUHRF1-high livers compared to controls, we found a trend towards hypomethylation (Figure 2B; 2E). We analyzed the methylation level of all CpGs that were fully methylated in control samples in the hUHRF1-high samples. This showed that 1.4% of these became fully unmethylated, while 7.7% reduced methylation levels to between 20-80%, suggestive of PMDs. Similarly, 1.1% of all CpGs that were unmethylated in controls became fully methylated in hUHRF1-high livers, while the remaining 3.2% increased methylation levels to 20- 80% methylation (Figure 2F). This indicates that only 2 days of hUHRF1 overexpression in hepatocytes can cause methylation disordering.

**Figure 2.**
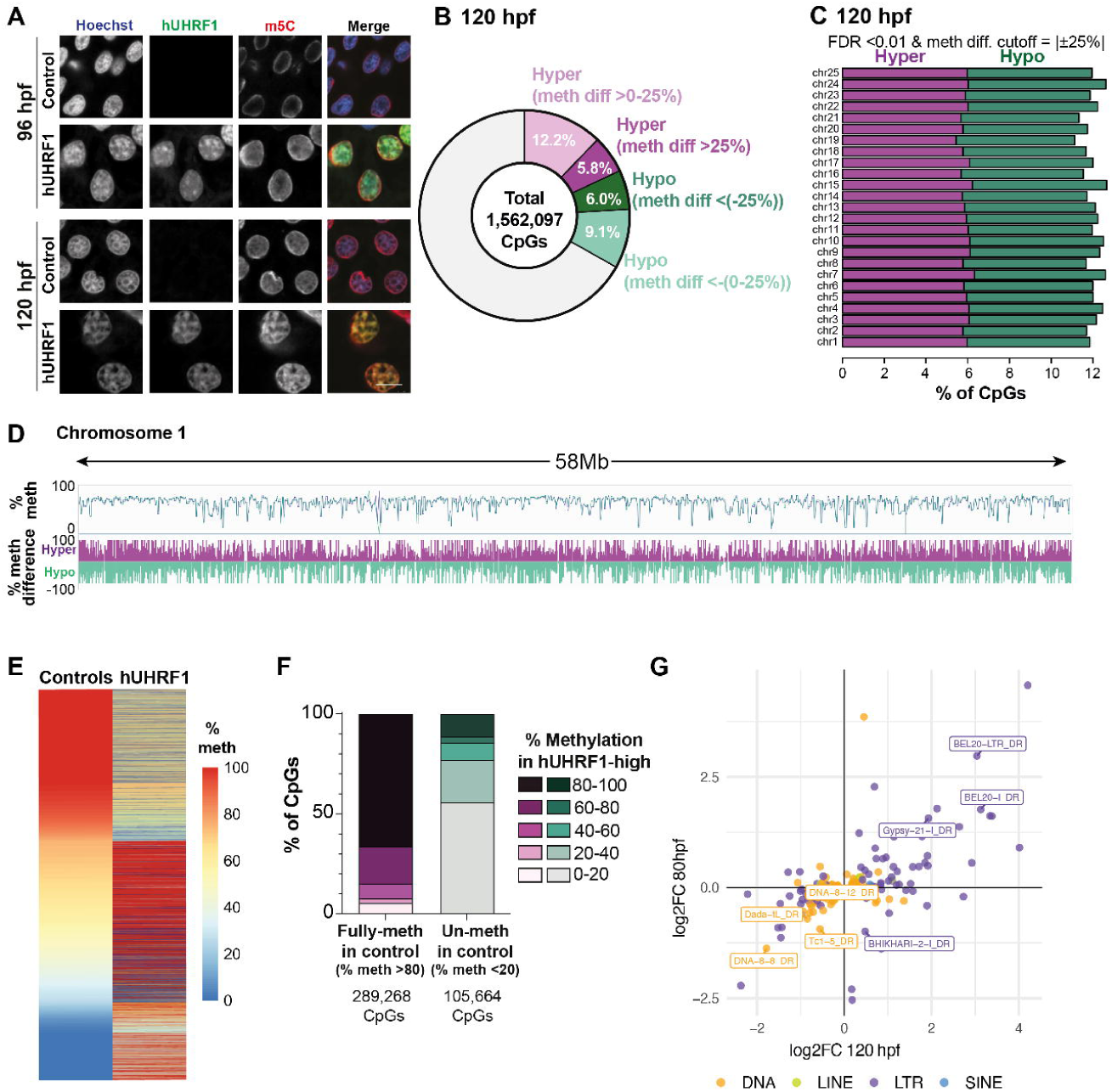
UHRF1 overexpression induces DNA methylome disordering and TE activation. **A.** Immunofluorescence assay of 5mC in hUHRF1 and control at 96 and 120 hpf. **B.** Pie chart of total number of CpGs covered by RRBS at 120 hpf zebrafish livers (1,562,097 CpGs). CpGs were categorized as: hyper-methylated >0-25% (methylation difference >0-25% & FDR <0.01; 190,576 CpGs), hyper-methylated >25% (methylation difference >25% & FDR <0.01; 90,986 CpGs), hypo-methylated <-(0-25%) (methylation difference <-(0-25%) & FDR <0.01; 142,626 CpGs) and hypo-methylated <-(25%) (methylation difference <-(25%) & FDR <0.01; 93,838 CpGs). **C.** Histogram of CpGs hyper-methylated (methylation difference >=25% & FDR <0.01) and hypo-methylated (methylation difference <=-(25%) & FDR <0.01) across all zebrafish chromosomes. **D.** Genome browser visualization of the chromosome 1. Track with percentage of methylation is an overlay of hUHRF1-EGFP (green) and control (blue) lineplots displaying per each condition the mean composite values calculated by default windowing function for all CpGs covered. Track representing differences in percentage of methylation is a bar chart displaying single not-combined values for all CpGs covered. **E.** Heatmap of differentially methylated CpGs (methylation difference >25% & <-(25%) & FDR <0.01) between hUHRF1 and controls. **F.** Stack bar plot of % of CpGs distribution in hUHRF1. CpGs were segregated in fully-methylated (% of methylation >80% & FDR <0.01) or un-methylated (% of methylation <20% & FDR <0.01) in controls showing a significant change in hUHRF1-EGFP and then stratified by % of methylation of hUHRF1-EGFP in five groups respectively. **G.** Crossplot of log2 Fold Change of repetitive elements in hUHRF1 and control at 80 and 120 hpf respectively. Names of TEs in the graph indicate TEs significantly changed at both and time points.

Heterochromatin loss and DNA methylation randomization can derepress TEs and this is proposed as a mechanism of mutagenesis in cancer^43^. To investigate the consequence of the epigenetic changes in hUHRF1-high livers, we examined TE expression by using bulk RNAseq data. As early as 80 hpf, many LTR retrotransposons were derepressed, and most persisted to 120 hpf (Figure 2G). This indicates that the epigenetic mechanisms that suppress TEs are lost as early as 80 hpf. Moreover, since TE retrotransposition generates DSBs, this is a potential mechanism of hUHRF1-induced DNA damage.

### UHRF1 causes p53-independent DNA replication arrest

We previously showed that loss of 1 copy of *tp53* reduces SA-β-gal staining, increases liver size and reduces time to tumor onset in hUHRF1-high animals^25^. We also showed that *atm* mutation enhanced the small liver phenotype and mortality caused by hUHRF1 overexpression^44^ indicating that the Atm-Tp53 pathways was functionally relevant to hUHRF1 induced changes. To investigate the mechanism by which hUHRF1 overexpression activates Tp53, we assessed the requirement for Atm and Atr in hUHRF1 induced senescence features. As expected, deletion of both copies of *tp53* in hUHRF1-high larvae completely abrogated SA-β-gal staining (Figure 2A). *atm* mutation also significantly decrease SA-β-galactosidase staining, while inhibition of Atr with VE-821 (Supplemental Figure S3) did not reduce SA-β-gal staining alone or in combination with *atm^-/-^* (Figure 3A). This indicates that both Atm and Tp53, but not Atr, are required for hUHRF1- induced SA-β-gal, and that Atr does not compensate for *atm* loss in this setting. Surprisingly, none of these interventions restored EdU incorporation in hUHRF1-high hepatocytes (Figure 2B). This indicates that hUHRF1 overexpression blocks proliferation directly, independent of Tp53.

**Figure 3.**
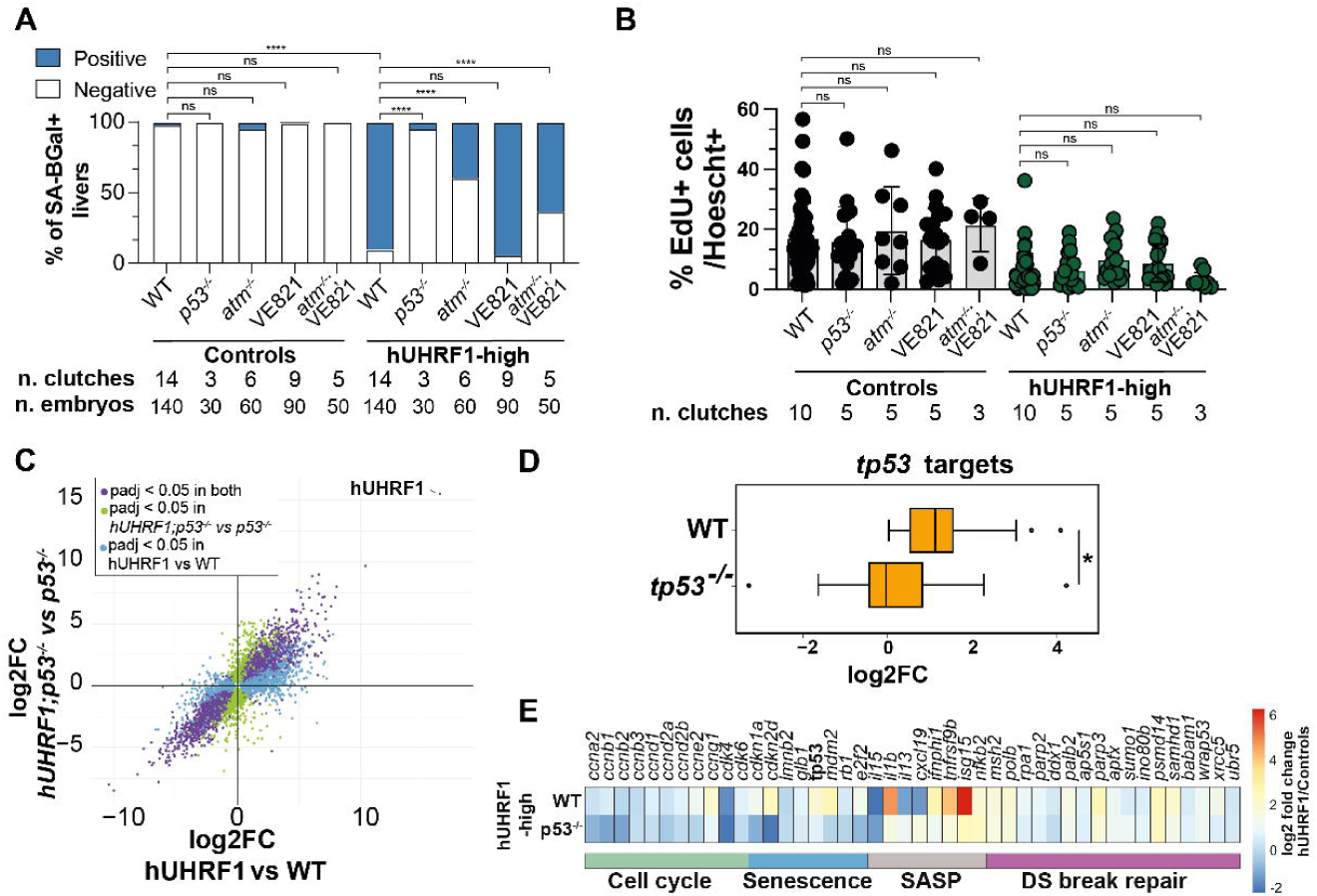
UHRF1 causes p53-independent DNA replication arrest. **A.** Stack bar plot of SA-ß-galactosidase positive livers at 120 hpf in hUHRF1 and control livers with deletion of tp53, atm, chemical inhibition of Atr (VE821) or combination of atm deletion and atr inhibition. Significance is measured by unpaired t-test. **** means p-values < 0.0001. **B.** EdU incorporation at 96 hpf in hUHRF1 and control livers with deletion of tp53, atm, chemical inhibition of Atr (VE821) or combination of atm deletion and atr inhibition. Significance is measured by unpaired t-test. **C.** Crossplot of log2 Fold Change of hUHRF1 compared controls and p53-/-; hUHRF1 compared p53-/- at 120 hpf. **D.** Box plot of log2 fold change of tp53 target genes^79^ upregulated in hUHRF1 compared wild type (WT, p-value adjusted < 0.05 & log2 fold change > 0) and tp53-/-; hUHRF1-EGFP compared to tp53-/- (tp53-/-). **E.** Heatmap of selected tp53 target genes involved in senescence in hUHRF1 compared to wild type (WT) and tp53-/-;hUHRF1 compared to tp53-/- (tp53-/-). For each experiment at least 3 biological replicates were performed.

To further investigate the hUHRF1-induced changes that require *tp53*, we carried out bulk RNAseq on pooled livers from WT and *tp53^-/-^* mutants with and without the hUHRF1-high allele (Figure S4; Table S1). hUHRF1 livers have over 7,000 DEGs at 120 hpf (padj < 0.05; Figure S4A- B). The majority of these were similarly deregulated in *tp53^-/-^* mutants (Figures 3C, Supplemental Figure S4C, D, E), with the exception of the canonical *tp53* target genes (Figure 3D). Importantly, genes involved in senescence and SASP which were upregulated in hUHRF1-high livers were downregulated when *tp53* was removed (Figure 3E; Table S1). This indicates that *atm* and *tp53* are required for specific senescence features caused by hUHRF1-high overexpression— *i.e.* SA- β-gal and the SASP. However, the cell cycle withdrawal induced by hUHRF1 is independent of these signals, suggesting that hUHRF1-high directly inhibits DNA replication, possibly by interaction with replisome factors.

### UHRF1 overexpression changes hepatocyte identity

To identify the cell specific gene expression changes caused by UHRF1 overexpression, we performed scRNAseq on pools of livers manually dissected from 120 hpf UHRF1-high and from control larvae where EGFP with a membrane targeting signal was expressed under the same hepatocyte promoter (*Tg(fabp10a:CAAX-EGFP);* hereafter called Controls; Figure 4A). After quality control, a combined dataset of 23,663 cells were segregated into 12 clusters that were assigned to different hepatic cell types according to expression of established cell identity markers^45,46^ (Figure 4B; Supplemental Figure S5A, Supplemental Table S4). There were 6 populations of hepatocytes and, as expected, together these constituted the majority of all cells (16,116 cells, 69.3% of total). The hUHRF1 and EGFP sequences were added to the danRer10 genome to allow transgene mapping, showing that in both control and hUHRF1 samples, the transgene was expressed primarily in the hepatocyte population (Figure 4C-D, S5B-C).

**Figure 4.**
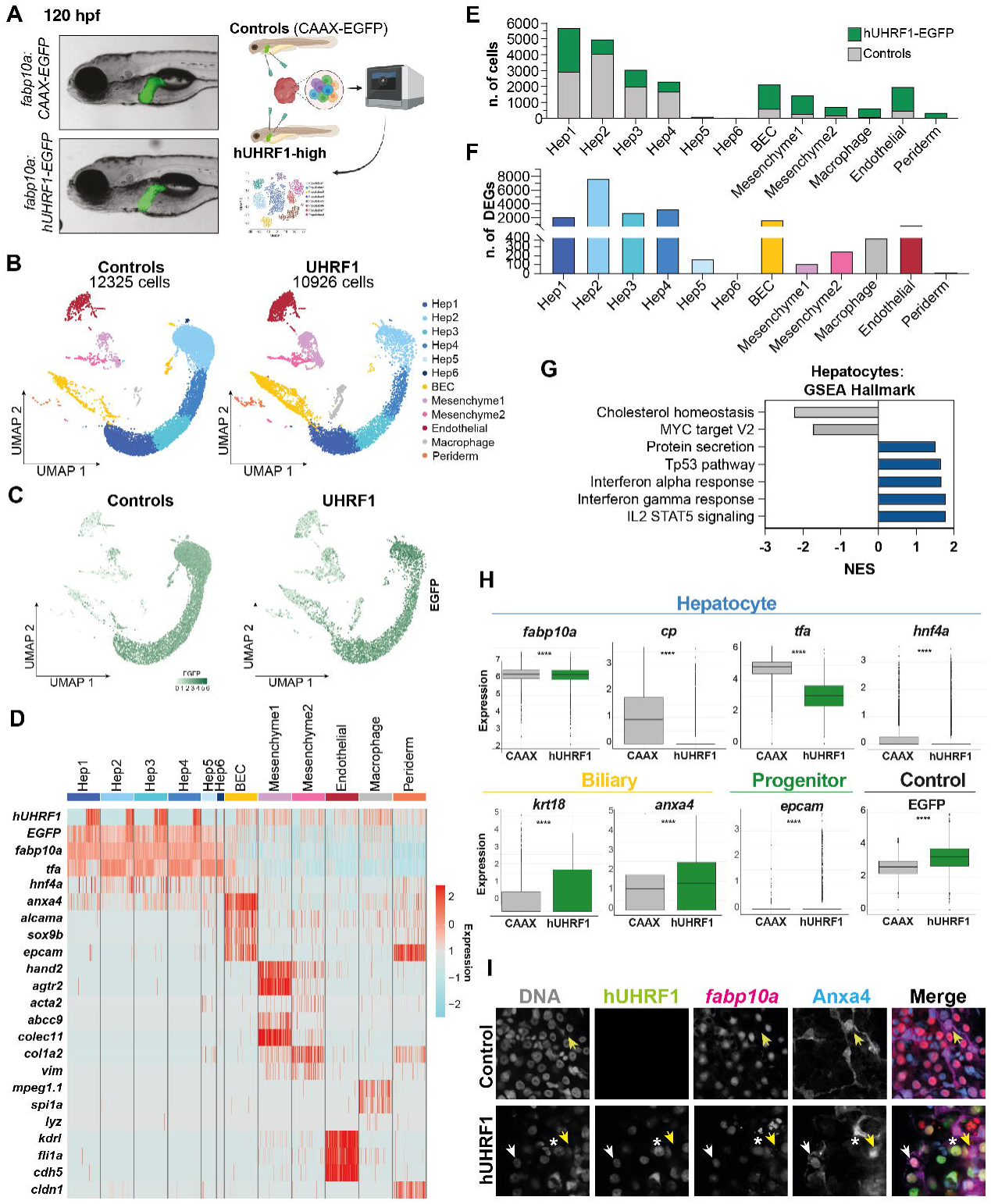
UHRF1 overexpression changes hepatocyte identity. **A.** Scheme of the approach and representative images of CAAX-EGFP (Controls) and hUHRF1- EGFP. **B.** UMAP plot of scRNAseq of hUHRF1 and control livers at 120 hpf, showing the different populations identified in livers. **C.** Feature plot of EGFP in hUHRF1 and control livers. **D.** Heatmap of key identity markers in different cell populations. **E**. Number of cells for each cluster in hUHRF1 and control livers. **F.** Number of differentially expressed genes (padj < 0.05) of each cluster in hUHRF1 compared to controls. **G.** GSEA of significant Hallmark pathways (FDR < 0.05) of differentially expressed genes (padj < 0.05) in all of Hepatocytes comparing hUHRF1 and CAAX. **H.** Box plot of expression values of hepatocyte, biliary epithelial cells and progenitor markers in hUHRF1 and control hepatocytes. **I.** Immunofluorescence of anxa4 in hUHRF1 and controls livers at 5 dpf. Star indicates hepatocyte positive for hUHRF1 and hepatocyte markers, yellow arrow indicates biliary cell positive for Anxa4 and white arrow indicates hepatocyte positive for hUHRF1, biliary and hepatocyte markers.

There was a striking shift in cell populations caused by overexpression of hUHRF1 (Figure 4E), with depletion of hepatocyte populations, especially Hep2 has high expression of the *hnf4a* transcription factor which drives hepatocyte identity (Figure 4D). Non-parenchymal cells represented only 12.7% of all cells in control samples, while they were 51% of all cells in hUHRF1- high samples (Figure 4E). Biliary epithelial cells (BEC), characterized by the expression of *alcama*, *sox9b*, *epcam* and keratin genes, represented the next largest population of cells in the combined dataset (2,216 cells; 9.1% of total), followed by endothelial and mesenchymal cells, which were divided into 2 populations(Figure 4B, D). The appearance of macrophages in the hUHRF1-high dataset (Figure 4B, E) confirms imaging data showing macrophage infiltration in the liver of hUHRF1 samples (Figure 1I).

The DEGs identified in the scRNAseq reflected and refined findings from bulk RNAseq data. There were thousands of DEGs in the hUHRF1-high samples, with the largest number of DEGs in the Hep2 population (Figure 4F). The DEGs in hepatocytes showed upregulation of genes representative of senescence, including induction of Tp53 and immune signaling, and downregulation of major hepatocyte functions, such as cholesterol metabolism (Figure 4G). The finding that there was a marked difference in the BEC population, with an increase in the number of cells and thousands of DEGs (Figure 4E-F), suggest that hUHRF1 overexpression both induces BEC expansion and changes their identity. To further investigate how hUHRF1 affects hepatic cell identity, we compared the expression of markers of hepatocyte, biliary and hepatic progenitor cell identity in the pooled hepatocyte populations from both samples. This showed that all markers of differentiated hepatocytes, including the master regulator of hepatic identity, *hnf4a*, were significantly down regulated in hUHRF1-high samples, while markers of biliary and progenitor identity are increased (Figure 4H). This suggests that hUHRF1 overexpression promotes stemness in hepatocytes and influences other cell types in the liver within 2 days of overexpression. This was confirmed by immunostaining livers for Anxa4, which identifies cells with a clear biliary morphology and is completely absent from all hepatocytes marked with the transgene (*fabp10a:nls-mCherry*) in controls (Figure 4I). In contrast, some Anxa4+ cells in hUHRF1-high samples express both hUHRF1-EGFP, and (*fabp10a:nls-mCherry*) (Figure 4I), indicating that hUHRF1 overexpression changes hepatocyte identity, promoting a progenitor like state.

A new finding from the scRNAseq was the marked expansion of mesenchymal cells in hUHRF1-high samples (Figure 4B, E). In controls, one population of mesenchymal cells expressed *hand2*, indicative of hepatic stellate cells, while another population was enriched for vimentin, and collagen, indicative of fibroblasts (Figure 4D). In hUHRF1 samples, the mesenchymal cells express *zeb2* and N-cadherin *(cdh2),* suggesting induction of EMT (Figure S6). Importantly, while some of these unique populations of BECs and mesenchymal cells expressed hUHRF1, the majority of them do not (Figure S5C, S6). Together, these data show that within 2 days of UHRF1 overexpression cell identity changes dramatically, including induction of EMT and stemness.

### UHRF1 induced heterogeneity and senescence escape leads to HCC

We determined whether hUHRF1 overexpression changes cell identity through a direct mechanism, requiring hUHRF1 expression, or whether the effect was mediated through epigenetic changes that were induced by hUHRF1 overexpression that could then persist even in the absence of UHRF1. To address this, we first established the pattern of hUHRF1-EGFP expression in hepatocytes by measuring the level of EGFP in individual hepatocytes from 80 hpf, when we first observed hUHRF1-mediated changes, through senescence (5 dpf) and pre-cancer stages (7, 10 dpf) and the first stage when cancer was observed (14 dpf)^25^. To determine if hUHRF1-EGFP was expressed in all hepatocytes, we compared EGFP expression to nls-mCherry driven by the same hepatocyte-specific promoter (*Tg(fabp10a:nls-mCherry)*; Figure 5A and S7A). After subtracting background fluorescence, we scored each nuclei for the presence of mCherry, EGFP or both. This showed that at 80 hpf, most mCherry-marked hepatocytes lack EGFP, but by 5 dpf, over 80% of hepatocytes expressing mCherry also express EGFP. However, by 10 dpf, nearly half of the mCherry positive nuclei no longer express EGFP (Figure 5B), indicating marked heterogeneity of hUHRF1-EGFP expression. To further evaluate this heterogeneity, we analyzed all nuclei expressing mCherry, and normalized the mCherry and EGFP fluorescence levels in each nuclei to their respective maximum and minimum values for each sample, thus allowing direct comparison of both fluorophores in each cell. As a control, we also assessed the expression of mCherry in samples which lacked hUHRF1-EGFP (Supplemental Figure S7B). We found that while there were only a few hepatocytes which expression levels of mCherry and EGFP are equivalent at 5 dpf, they progressively become mismatched, so by 7 dpf, many of the mCherry expressing cells have low EGFP and *vice versa* (Figure 5C and Supplemental Figure S7A-B). Importantly, this data reveals a highly heterogenous expression pattern of EGFP hepatocytes. *In situ* hybridization for human UHRF1 showed that this heterogeneity is present at mRNA level (Supplemental Figure S7C) suggesting that some hepatocytes downregulate hUHRF1-EGFP expression, resulting in heterogeneity of this oncogene during hepatocarcinogenesis. It is also possible that since UHRF1 overexpression induces a stemlike state it reduces expression of hepatocyte specific genes, including the *fapb10a* transgene promoter (Figure 4H).

**Figure 5.**
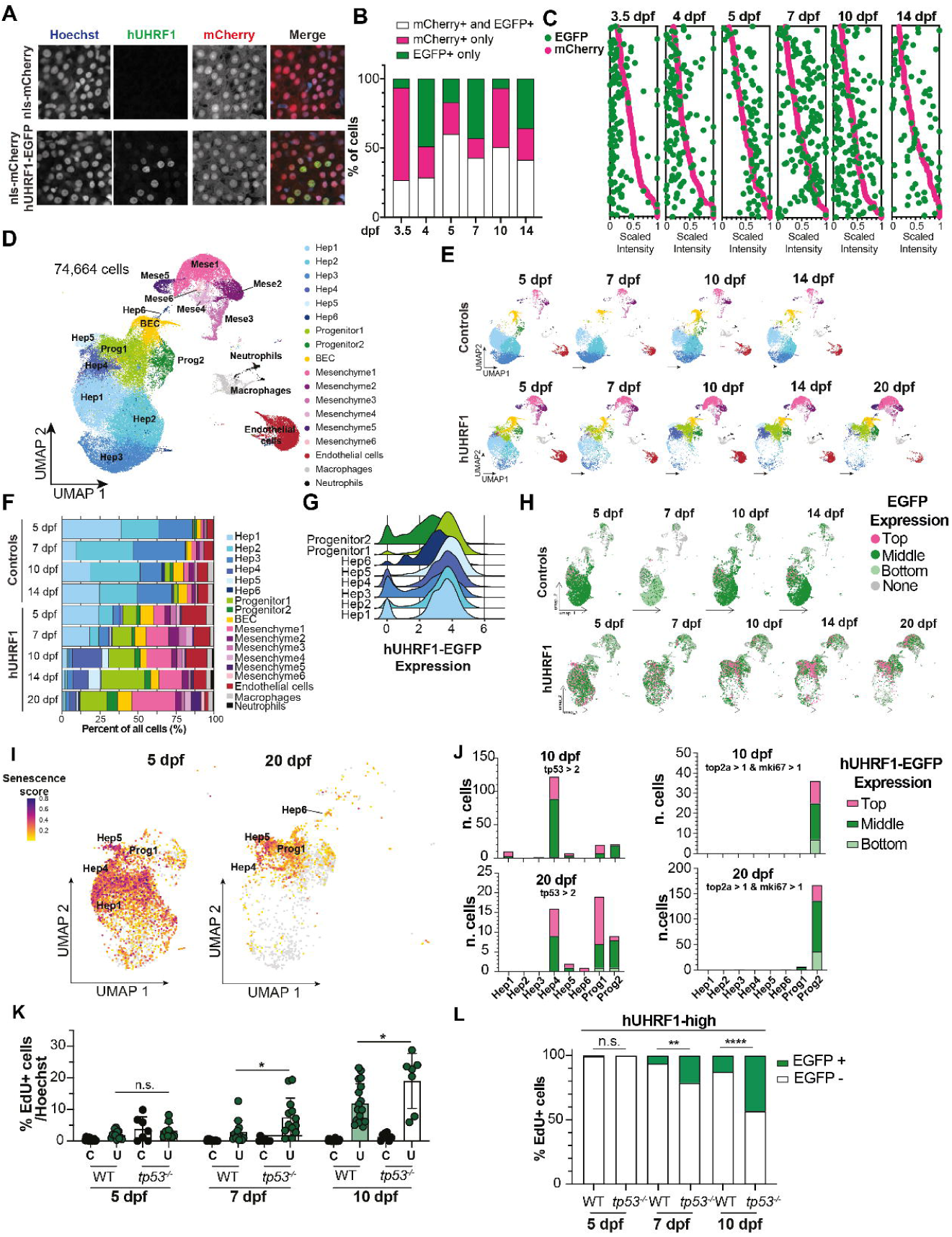
UHRF1 induced heterogeneity and senescence escape leads to HCC. **A.** Representative images of hUHRF1;nls-mCherry and nls-mCherry control livers at 5 dpf. **B.** Quantification of cells positive for EGFP and nls-mCherry, EGFP only and nls-mCherry only in hUHRF1;nls-mCherry livers at different time points. **C.** Quantification of Scaled Fluorescent Intensity normalized on maximum intensity for each channel for each liver of EGFP and nls-mCherry in hUHRF1;nls-mCherry livers across time points. **D.** UMAP plot of scRNAseq of hUHRF1 and control livers combined at 5, 7, 10, 14 and 20 dpf showing the different populations identified in the liver. **E.** UMAP plot of scRNAseq of hUHRF1 and control livers across time points. **F.** Stack bar of different populations showed in D for hUHRF1 and controls at each time point. **G.** Ridge plot of hUHRF1 in hepatocytes and progenitors populations combining all time points. **H.** UMAP plot of scRNAseq of EGFP levels in hUHRF1 and control livers across time points divided in top (EGFP > 3.71), middle (EGFP < 3.71 & EGFP > 1.78), bottom (EGFP < 1.78 & EGFP > 0.01) and no expressing EGFP (EGFP < 0.01) based on Gaussian distribution of EGFP in hUHRF1 and control livers. **I.** Feature plot of senescence features at 5 dpf and 20 dpf defined as top 15 differentially expressed genes in senescent hepatocytes at 5 dpf. **L.** Stack bar of number of cells for each population at 10 dpf and 20 dpf that are senescent (tp53 > 2) or proliferative (mki67 >1 & top2a >1) divided in top, middle and bottom expression of hUHRF1. **M.** EdU incorporation assay in hUHRF1-EGFP, WT control, tp53^-/-^;hUHRF1 and tp53^-/-^ livers at 5, 7, 10 dpf. Significance is measured by unpaired t-test. * means p-values < 0.05. **N.** Stack bar of percentage of EdU positive cells in hUHRF1 livers divided in hUHRF1 positive and hUHRF1 negative cells. Significance is measured by Chi-square test, ** indicates p-value < 0.01 and **** indicates p-value < 0.0001.

To further investigate the mechanism of epigenetic driven transformation, we carried out scRNAseq on cells from dissected livers from hUHRF1-high and *Tg(fabp10a:CAAX-EGFP)* during the time points covering the complete trajectory of tumorigenesis: senescence (5 dpf), preneoplastic (7-10 dpf) and cancer (14-20 dpf). Combined, nearly 75,000 cells were sequenced, obtained from pools of over 2,000 dissected livers per genotype. The unified dataset was segregated into distinct cell populations based on top differentially expressed markers (Supplemental Figure S8A, Supplemental Table S5) and previously published markers for zebrafish liver^45,46^. After cleaning the dataset to remove minor populations of non-hepatic cells, they remained 6 populations of hepatocytes, BECs, 6 populations of mesenchymal cells, endothelial and immune cells, representing neutrophils and macrophages (Figure 5D).

Cells in controls and hUHRF1-high samples were segregated to examine how these populations changed over time (Figure 5E-F). While hepatocyte populations were relatively constant in controls, hepatocyte clusters which express the majority of the hepatic function genes are progressively eliminated in the hUHRF1 samples. Interestingly, several unique cell populations emerge during the course of tumorigenesis, including hepatocyte clusters (Hep 4 and 5) and progenitors (Figure 5E-F and Supplemental Figure S8B).

This analysis identified substantial cellular heterogeneity in hUHRF1-high samples at all time points, which became most apparent at 20 dpf, with hepatocytes making up the minority of cell types, and progenitor and mesenchymal cells predominating (Figure 5F and Supplemental Figure S8B). Since the majority of hUHRF1-high fish have HCC at 20 dpf, we conclude that the populations of cells identified at this time point represent both malignant as well as tumor-supporting cells, including immune and mesenchymal cells which could serve as cancer fibroblasts. By combining all cells from all time points, we were able to define populations of cells that could not be identified when analyzing only the 5 dpf dataset, revealing progenitor cells were present as early as 5 dpf, and expanded to become the predominant cell type in the cancer stages (Figure 5E-F and Supplemental Figure S8B). This suggests that the identity of the cells that form HCC is established early after hUHRF1 overexpression.

We hypothesized that persistent and high levels of hUHRF1 expression would sustain senescent cells, and that the tumor cells would emerge from those that had downregulated or silenced hUHRF1 expression, similar to findings with Ras overexpression in hepatocarcinogenesis^3^. To test this, we asked whether EGFP expression levels correlated with a distinct cell population. We found highest levels in Hep5 and lowest levels in Hep3 at all time points (Supplementary Figure S9). We next examined the range of EGFP expression in combined hepatocyte and progenitor populations from all time points which showed that all cells in the Hep5, Hep6 and Progenitor1 populations maintained hUHRF1-EGFP expression levels over time, while there was a large range of expression in Hep1, Hep3, Hep4 and Progenitor2 populations, with Progenitor2 cells having the most heterogeneous levels (Figure 5G). This indicates that those cells observed by imaging that have heterogenous hUHRF1-EGFP expression represent distinct hepatocyte populations which can be differentiated based on their gene expression profile. To further identify the features of cells with different expression levels, we calculated the Gaussian distribution of EGFP in all cells at all time points of both hUHRF1-high and control samples and categorized cells based on expression quartiles into top, middle, low and no expression levels (Supplemental Figure S9B). UMAP analysis of cells in control samples show that the highest-expressing cells (top and middle levels) were evenly distributed across all hepatocyte populations. In contrast, in hUHRF1-high samples, the highest-expressing cells were spread throughout hepatocyte populations at 5 dpf, but then became concentrated in specific hepatocyte populations (Hep4/5/6) at later stages; there were also a cluster of the highest expressing cells in the progenitor populations during cancer emergence (Figure 5H). Indeed, a senescence gene signature shows shift from Hep1-Hep4 populations at 5 dpf to Hep5-6 cells at 20 dpf (Figure 5I), indicating that a small population senescent cells with very high hUHRF1-EGFP levels persist during hepatocarcinogenesis.

To further test the hypothesis that cells with high levels of hUHRF1-EGFP are senescent and non-proliferative, we segregated all hUHRF1-EGFP expressing cells at 10 and 20 dpf and then categorized them as positive for expression of a marker of senescence (*tp53*) or proliferation (*ki67* and *top2a*). This shows that cells with high hUHRF1-EGFP – i.e. Hep4 and Hep5 populations - were positive for *tp53* and negative for *top2a* and *mki67* (Figure 5J) while Progenitor2 cells, which have a range of hUHRF1-EGFP expression levels were positive for markers of proliferation (Figure 5J). Together, these data indicate that tumor heterogeneity in this model has persistent senescent cells with high hUHRF1-EGFP levels and high *tp53*, intermingled with proliferating cells that have a range of hUHRF1-EGFP levels and downregulated *tp53*, suggesting these have escaped senescence and are malignant.

To examine this further, we examined proliferation in hUHRF1-high and control samples during the time course of tumorigenesis. We previously showed that 46% of hUHRF1-high animals had HCC by 15 dpf^25^, indicating that the proliferative block induced by hUHRF1 was bypassed before this time. EdU incorporation showed increased proliferation in hUHRF1-high samples as early as 7 dpf and progressed to 10 dpf, when 11.9 % of cells in hUHRF1-high livers incorporated EdU (Figure 5K). We tested the hypothesis cells expressing high levels of hUHRF1 were senescent and therefore could only proliferate if *tp53* signaling was abolished by removing *tp53* and examining proliferation. The number of proliferating cells were significantly increased in hUHRF1-high; *tp53^-/-^* samples at 7 and 10 dpf to 7.8% and 18.2% in *hUHRF1-high;tp53^-/-^* samples, respectively (Figure 5K). Importantly, loss of *tp53* increased the number of proliferating hUHRF1- EGFP expressing cells. This indicates that some hepatocytes were able to escape hUHRF1 induced senescence during tumorigenesis, and that *tp53* is a gatekeeper to senescence escape.

Together, these data suggests that Progenitor2 cells are the proliferative cells that give rise to HCC, suggesting that these cells escape senescence.

### UHRF1-induced cancer cells originate from damaged progenitors

Analysis of scRNAseq datasets of hUHRF1-high samples identified two distinct populations of cells that have stemlike features: Progenitor 1 and Progenitor 2 (Supplemental Figure S10). These populations have low expression of hepatocyte markers compared with hepatocytes that express normal levels of genes regulating hepatic function (Hep3) and with senescent cells that express high levels of hUHRF1-EGFP and *tp53* (Hep5) (Figure 1A). These progenitor populations also express high expression of biliary markers (*anxa4*) and stem cell markers including *epcam*, *prom1a* and others (Figure 6A and Supplemental Figure S10). Markers of proliferating cells, such as *mki67* were predominantly detected in Progenitor2 (Figure 6A). Together this defines a novel population of stemlike cells which we speculate represent the cancer cells that have escaped senescence.

**Figure 6.**
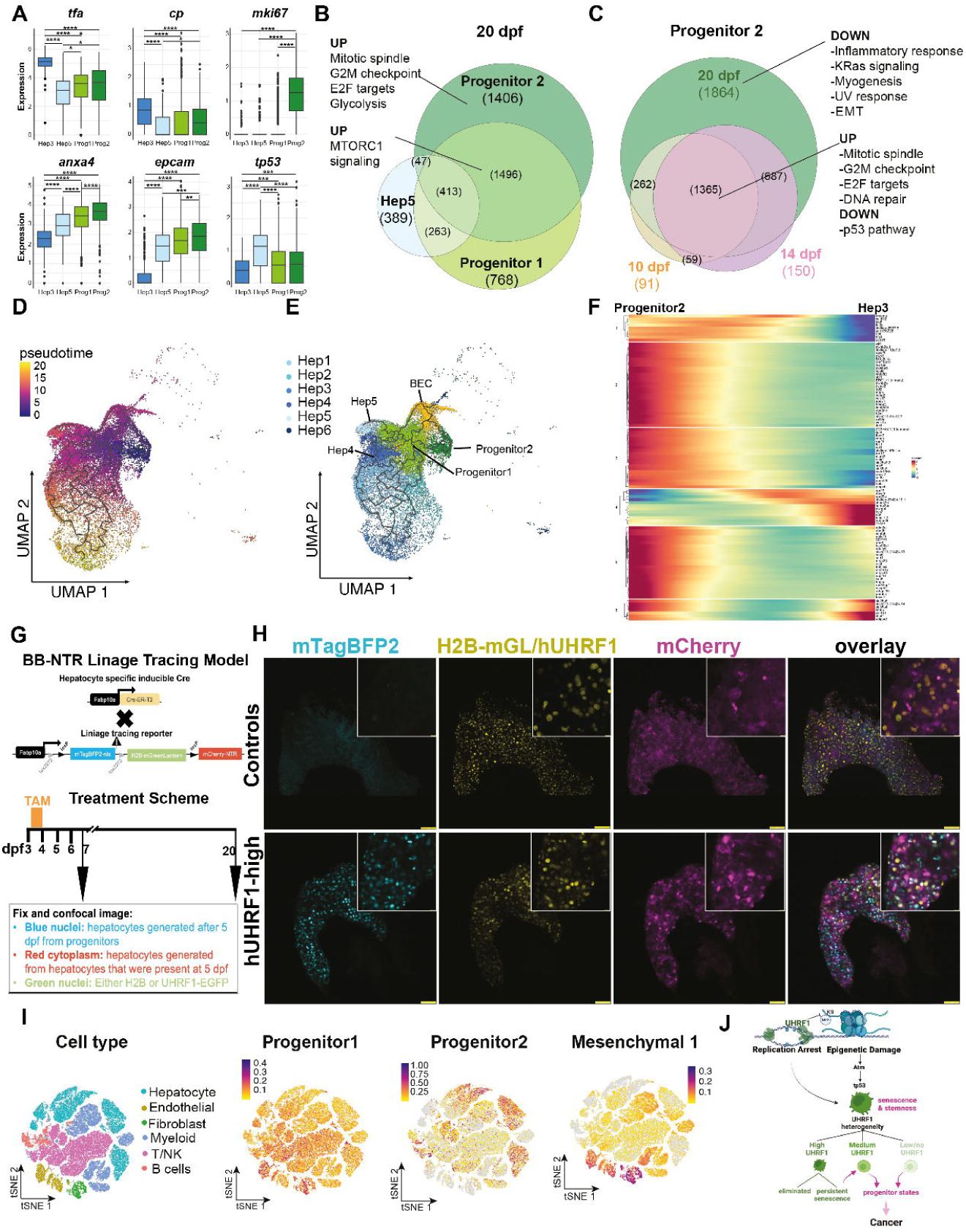
UHRF1-induced cancer cells originate from damaged progenitors. **A.** Box plot of hepatocytes, biliary epithelial cells, and proliferation markers in Hep3, Hep6, Progenitor 1 and Progenitor 2 at 20 dpf in hUHRF1. Significance is measured by unpaired t-test. * means p-value < 0.05, ** means p-value < 0.01, *** means p-value < 0.001, **** means p-value < 0.0001. **B.** Venn diagram of differentially expressed genes (padj < 0.05) at 20 dpf in Progenitor 2, Progenitor 1 and Hep5 calculated on all cells at 20 dpf. GSEA of upregulated and downregulated Hallmark pathways (padj < 0.05) of genes in each group of the Venn diagram. **C.** Venn diagram of differentially expressed genes of Progenitor 2 across time points calculated on expressed genes in Progenitor 2 compared to all other cells present at the same time point. **D.** Pseudotime projection of hepatocytes, biliary epithelial cells, progenitors in hUHRF1 livers including 5, 7, 10, 14, 20 dpf. **E.** UMAP on hepatocytes, biliary epithelial cells, progenitors in hUHRF1 livers including 5, 7, 10, 14, 20 dpf used to perform pseudotime projection. **F.** Expression heatmap of top significant 100 genes based on q-values driving underpinning pseudotime trajectory showing kinetic trends of common DEGs shared between Progenitor 2 and Hep3 plotted in pseudotime. **G.** Scheme of lineage tracing with fabp10a:BB-NTR; fabp10a:CreER and treatment scheme. **H.** Representative images of fabp10a:BB-NTR; fabp10a:CreER (Controls) and hUHRF1; fabp10a:BB-NTR; fabp10a:CreER (hUHRF1) at 20 dpf. **I.** tSNE plot of human HCC showing different cell types across 10 HCC patients and feature plots of signature of progenitor 1, progenitor 2 and mesenchymal 1 identified in zebrafish livers. **J.** Model of the relation between UHRF1 levels, senescence, stem-like state and cancer formation.

To further investigate the identity and features of these cell populations, we identified the DEGs in both progenitor populations compared to the persistent senescent cells (Hep5) at 20 dpf, when malignant cells are present in nearly every liver. GSEA showed that the Progenitor2 cells were enriched for proliferation related pathways, including glycolysis, a key metabolic pathway utilized by malignant cells facilitating their rapid growth and survival^47^. Both Progenitor1 and Progenitor2 cells have upregulation of Mtorc signaling, key cancer pathway (Figure 6B). To identify how Progenitor2 cells evolve over the course of tumorigenesis we evaluated the DEGs in this population at 10, 14 and 20 dpf compared to all other populations in the hUHRF1-high samples. This showed that the transcriptome was largely similar in the pre-cancer timepoints, and there were 1,365 DEGs common to all time points were enriched for cell proliferation and Tp53 signaling was downregulated (Figure 6C). However, at 20 dpf, there were nearly 2,000 DEGs, characterized by downregulation of inflammatory pathways and UV damage response, which is characteristic of double stranded breaks, suggesting that these have downregulated the persistent DDR and SASP that characterizes senescent cells.

Overall, these data demonstrate that Progenitor2 cells exhibit key features of cancer cells: glycolysis, stemness, proliferation and loss of Tp53 mediated tumor suppression. We hypothesized that these cells derived from previously senescent hepatocytes that escaped *tp53* activation by lowering hUHRF1-EGFP. To test this, we generated a pseudotemporal trajectory of Hep1-6, Progenitor1-2 and BECs from hUHRF1-high samples at all time points. We designated Progenitor2 as the root to trace the origin of these presumed cancer and found that Progenitor2 and Hep3 are the most distantly related populations. Conversely, there is a direct, unbranched linage from Progenitor1 (Figure 6D-E), suggesting that Progenitor1 serves as precursor population. From Progenitor1, there is a unidirectional trajectory from BEC cells, suggesting that some of these have a linear progression from this cell type, and, importantly there is also a liniar trajectory from Hep4 and Hep5, the persitantly senescent cells (Figure 6D-E). Thus, Progenitor1 are likely the intermeidate cell stage that has escaped senescnece, reduced hUHRF1-EGFP levels but has not yet fully aquired the metabolic and proliferative capacities of the Progenitor2 cancer cells.

We tested the hypothesis that the Progenitor2 cancer cells originate from previously senescent cells using a linage tracing strategy in which hepatocyte specific promoter (*fabpp10a*) drives the expression of a cassette containing mTagBFP2 between loxP sites followed by H2B- mGL and cytoplasmic mCherry. These were crossed to a transgenic line with hepatocyte specific inducible Cre (*Tg(fabp10a:CreERT2,cryaa:ECFP*); Figure 6G) and were treated with tamoxifen at a time when senescence induction is underway (3.5-4 dpf) to mark all senescent cells and then evaluated the cell populations at 20 dpf. In controls that are not treated with tamoxifen, all cells are BFP positive at all time points, and tamoxifen effectively removes the mTagBFP2 signal and induces H2B-mGL and mCherry-NTR (Supplemental Figure S11A-B). If cells are generated new from BECs after 4 dpf, they will express mTagBFP2, and if they are derived from senescent hepatocytes, they will be marked by mCherry. As expected, controls treated with tamoxifen showed that all cells in the 20 dpf liver were derived from hepatocytes (i.e. were all mCherry or GFP positive Figure 6H). Since the H2B-mGL and hUHRF1-EGFP have the same signal, we relied on mCherry expression as a marker of hepatocyte linage. At 20 dpf, hUHRF1-high livers are riddled with HCC, and the cells in these samples were predominantly mCherry positive (Figure 6H) intermingled with BFP positive cells. This is consistent with previous studies showing that a small population of progenitor cells contribute to hepatocytes hUHRF1-high livers^48^.

This *in vivo* data, combined with the scRNA-seq analysis, demonstrates that UHRF1 overexpression drives tumor formation by inducing cancer cells derived from two sources: previously senescent hepatocytes present at 4 dpf and new hepatocytes generated as a regenerative response to hepatocyte loss and impaired hepatic function. The resulting tumor is heterogeneous, displaying variability in cell identity and in the levels of hUHRF1 expression across distinct cell populations. We conclude that UHRF1 promotes the emergence of diverse progenitor-like cancer cells, which are proliferative and arise from epigenetically altered hepatocytes that have escaped senescence.

### UHRF1-induced progenitor signatures demark human HCC cell populations

UHRF1 is upregulated in most human solid tumors, and it has been considered a tumor biomarker^49^. We postulate that in many cases this is due to a bystander effect, whereby UHRF1 reflects the proliferative index of these tumors, but in a subset of tumors characterized by the Progenitor2 expression signature, it serves as a driver. To test this, we analyzed the expression of the progenitor cell signatures identified from hUHRF1-high zebrafish tumors in publicly available scRNAseq datasets obtained from 10 HCC patients^50^. Strikingly, the Progenitor1 signature was elevated across all cell types in these samples (Figure 6I), indicating lack of specificity of the genes that define these cells. In contrast the Progenitor2 signature highlighted a distinct population of cells, corresponding to a subset of HCCs (Supplemental Figure S12). This analysis also showed that the fibroblasts in the human HCC exhibited high expression of a mesenchymal-like cell signature found in the mesenchymal cells from hUHRF1-high samples (Figure 6I), suggesting that this population of mesenchymal cells are cancer fibroblasts. Together, this suggests that UHRF1 induces a gene expression signature defining a subset of human HCCs, suggesting this as a driver of this subset of human HCCs.

## Discussion

Epigenetic alterations are well-established hallmarks of cancer; however, their mechanistic contributions to oncogenesis remain poorly defined. Identifying tumor subtypes amenable to therapeutic targeting of epigenetic mechanisms has driven efforts to develop such therapies, however, this has not yet translated to major clinical impact. This may be attributed to the fact that epigenetic changes can be accompanied by or causative of genomic changes, which are irreversible even if the epigenome is restored. Therefore, it is critical to identify the tumor subtypes in which epigenetic changes are the primary driver in the absence of genomic instability. The encouraging finding that a short depletion of the polycomb complex in *Drosophila* can cause cancer in the absence of any genomic changes^23^ shows that epigenetic changes alone provide a sufficient oncogenic stimuli. However, it is not yet clear whether this mechanism occurs in other models or in humans. We examined this in a model of hUHRF1 overexpression in zebrafish hepatocytes which induces senescence followed by a high incidence of HCC within 2 weeks of transgene induction^25^.

Here we show that randomized DNA methylation and transposon activation occurs within 1 day of hUHRF1 overexpression in this model and is accompanied by DNA damage, triggering Atm and Tp53 to induce senescence characterized by SASP and immune cell recruitment. We also provide evidence that high levels of hUHRF1 block DNA replication directly and that cells with very high expression are eliminated over time, as found for hepatocytes expressing high levels of Ras^3,6^. This zebrafish model identified a unique mechanism of hepatocarcinogenesis driven by epigenetic rearrangement due to hUHRF1 overexpression. Importantly, these changes occur rapidly and first induce senescence due to DNA damage, Atm and Tp53 activation. The majority of these damaged cells are eliminated, but some persist and escape to generate heterogenous tumors that include cancer cells with stem-like features, persistent senescent cells, cancer fibroblasts and immune cells. Since the UHRF1 driven tumors are caused by loss of a repressive epigenome and genomic instability, cancers resulting from UHRF1 overexpression may not be amenable to treatment by therapies aimed at restoring the epigenome.

How the epigenome is repatterned in cancer is not known. One possibility is that DNA methylation changes occur during oncogene induced senescence^12–14^ due to deregulation of epigenetic writers and readers; when these cells escape senescence, the epigenetic patterns that are beneficial to cancer cells are maintained during tumor evolution. We propose that in a subset of tumors, UHRF1 overexpression is the cause of methylome repatterning. This phenomenon likely occurs in settings where UHRF1 is transiently overexpressed in premalignant cells, and once DNA methylation patterns are altered, they can be stably propagated during cell division, providing UHRF1 is present during replication. Identifying such cases is challenging, as even brief, high levels of UHRF1 expression may suffice to induce these epigenetic changes. We propose that most cells with epigenetic damage and high UHRF1 levels undergo senescence and are eliminated. A recent studies in a Ras model of hepatocarcinogenesis elucidated the importance of oncogene titration, with high Ras levels inducing senescence and immune clearance, while moderate Ras overexpression promotes an immune-evasive, stem-like state capable of escaping senescence and HCC^3^. This phenomenon parallels our findings with UHRF1: high UHRF1 levels induce senescence, which acts as an effective tumor-suppressive mechanism. However, a subset of these senescent cells, through an unknown mechanism, reduces UHRF1 levels while retaining epigenetic alterations and DNA damage. These damaged, progenitor-like cells persist and, with the loss of *TP53*, can re-enter the cell cycle, undergo malignant transformation, and contribute to tumor heterogeneity. This underscores the dual role of UHRF1- induced senescence in both suppressing tumor development and initiating cancer.

We observed a high level of tumor heterogeneity caused by UHRF1. This is a common feature of HCC, which has significant inter and intra tumor heterogeneity^51–54^, and is a linchpin of treatment as only a few resistant cells can cause therapy resistance^2^. Importantly, the intratumoral heterogeneity in HCC is reflected in DNA methylation heterogenetity^51,52,55^, suggesting that there is an initiating event that randomizes the methylome, and then patterns that confer selective advantage during tumor evolution are amplified. The methylome in most HCCs is not only heterogeneous, but the tumor also exhibits significant cellular diversity, including immune cells, cancer-associated fibroblasts, stromal cells, and cancer cells spanning a spectrum from progenitor-like to fully differentiated states. While gene mutations contribute to this diversity, alterations in the epigenetic landscape provide a rapid mechanism to change the genes that drive cell identity. We propose that the epigenetic changes caused by UHRF1 overexpression as a means to achieve heterogeneity in both the methylome and cell identity during HCC evolution.

These findings have significant implications for human cancers, as UHRF1 is not only overexpressed in hepatocellular carcinoma (HCC) but also in the majority of other cancer types, where overexpression is strongly associated with tumor progression and poor prognosis^56^. Interestingly, however, UHRF1 is expressed in most tumors in a salt and pepper pattern^25,57^, indicating that not all cancer cells require it’s expression. One possible explaination for this finding is that the elevated levels of UHRF1 in tumors may result from its upregulation during S phase^58,59^ and degraded during mitosis^60^ reflecting a high proliferative index and heterogeneous expression across tumor cells. As an essential component of the DNA replication machinery, UHRF1 in these cases is critical for cancer cell proliferation. This is supported by studies showing that targeting UHRF1 in cancer cells activates tumor-suppressive processes such as apoptosis and senescence^30,49,56,61,62^, similar to the findings of depleting Uhrf1 in highly proliferative cells during development^63–65^. This is further supported by the report that Uhrf1 deficient hepatocytes are refractory to chemical and genetically induced HCC^57^. Thus, targeting UHRF1 in these cases may effectively block cancer cell proliferation.

Our findings define a distinct cancer subtype in which hUHRF1 acts as a critical oncogenic driver. In this context, UHRF1 overexpression induces cancer-associated methylation randomization, *TP53*-dependent senescence, and a proliferation block. However, if these cells evade clearance and subsequently downregulate UHRF1 and *TP53*, they overcome the proliferation block and senescence. This process facilitates profound cell identity changes, drives genomic instability, and creates a permissive environment for tumor evolution.

## Acknowledgments

This work was supported by NYUAD Faculty Research Fund (AD188), NIH (5R01DK080789-11) to KCS and Tamkeen under the NYU Abu Dhabi Research Institute Award to the NYUAD Center for Genomics and Systems Biology (ADHPG-CGSB). All imaging was carried out in the NYUAD Core Technology Platform Imaging facility with expert support by Rashid Razgui. RNAseq bulk libraries were performed by NYUAD Core Technology Platform Imaging with expert help of Marc Arnoux and Mehar Sultan. Bioinformatics assistance was provided by the NYUAD Bioinformatics Core at New York University Abu Dhabi. We would like to thank Nizar Drou, Muhammad Arshad and Giuseppe-Antonio Saldi for assistance with deep sequencing, DNA methylation analysis, TE analysis and scRNAseq trajectory.

## Author contributions

EM and KCS conceptualized the project, EM, FM, BM, CC and SR developed the methodology, EM, FM, TR and CC carried out the analysis, EM, FM, YA, BM, TR, CC and SR carried out the investigation, EM, FM, CC and TR created the visualization, KCS provided resources, supervision, project administration, wrote the first draft and EM and KCS reviewed and edited the manuscript.

## Supplemental information

Supplemental Figure S1. hUHRF1 phenotype does not depends on integration site.

Supplemental Figure S2. RNAseq of 80 hpf hUHRF1 compared Controls

Supplemental Figure S3. Validation of Atr inhibition.

Supplemental Figure S4. RNAseq of 120 hpf hUHRF1 compared Controls and tp53-/-; hUHRF1 compared to tp53-/-.

Supplemental Figure S5. scRNAseq of hUHRF1 and Controls at 5 dpf.

Supplemental Figure S6. EMT markers in scRNAseq.

Supplemental Figure S7. hUHRF1 heterogeneity in hUHRF1 livers.

Supplemental Figure S8. scRNAseq of hUHRF1 and Controls at 5, 7, 10, 14 and 20 dpf.

Supplemental Figure S9. EGFP across population in scRNAseq.

Supplemental Figure S10. Feature Plot of genes upregulated in stem-like cells in the liver.

Supplemental Figure S11. Lineage-traced larvae at 6 dpf.

Supplemental table S1. Bulk RNAseq at 80 hpf. *(Available upon request)*

Supplemental table S2. Bulk RNAseq at 120 hpf. *(Available upon request)*

Supplemental table S3. Bulk RNAseq at 120 hpf in tp53-/-. *(Available upon request)*

Supplemental table S4. scRNAseq 5 dpf. *(Available upon request)*

Supplemental table S5. scRNAseq 5, 7, 10, 14, 20 dpf. *(Available upon request)*

## Materials and Methods

### Generation of zebrafish line

*Tg(mpeg1.1:dsRed2)* fish expressing dsRed2 only in macrophages was generated by PCR amplification of dsRed2^66^. BamH1 (New England Biolabs) and MfeI (New England Biolabs) restriction enzyme sites were inserted with PCR (Q5 Taq, New England Biolabs) in the forward and reverse primer respectively. A vector containing tol2 sites and mpeg1.1:Dendra2 cassette (Addgene: #51462) was digested with the same restriction enzymes. After ligation, transformation in DH5alpha (Thermo Fisher Scientific) and sequencing of positive colonies, 1 nl of 40 ng/μl of plasmid was injected with 80 ng/μl ng of Tol2 transposase mRNA into 1-2 cell stage embryos. Once embryos reached sexual maturity, they were crossed to wild type to generate F1 and selected for bright expression of dsRed2 into macrophages.

*Tg(fabp10a:lox2272-loxp-nls-mTagBFP2-stop-lox2272-H2B-mGL-stop-loxp-mCherry-NTR; cryaa:mCherry)^ulb^*^33^ (hereafter referred as fabp10a:BB-NTR, transgenesis marker “red eye”) and *Tg(fabp10a:CreER; cryaa:CFP)^ulb^*^34^ (hereafter referred as fabp10a:CreER, transgenesis marker “green eye”) was injected into 1-2 cell stage wilt type embryos with meganuclease to generate trangenic line. Once embryos reached sexual maturity they were crossed to wild type to generate F1 and selected for the presence of the cassette.

### Zebrafish Husbandry

Zebrafish (*Danio rerio*) husbandry and care was conducted according to New York University Abu Dhabi (NYUAD) for Animal Care and Use Committee (IACUC) Committee (protocol number 22- 0003A2). Adult fish were raised on a 14:10 h light: dark cycle at 28 °C and fed 2 times a day with brineshrimps and once with solid food. Larvae after 5 dpf were fed two times daily with paramecia untill 12 dpf and then with brine shrimp until 20 dpf. *Tg(fabp10a:hsa.UHRF1-EGFP^mss1^)^high^*^25^ called hUHRF1, Tg(*fabp10a:CAAX-EGFP)*^67^, Tg(*fabp10a:nls-mCherry)*^25^, Tg(*mpeg1.1:dsRed2)* were crossed to wild type (WT) zebrafish adults or to transgenic line to generate hUHRF1^+/−^ zebrafish embryos. *tp53^−/−^* ^68^ and *atm^−/−^*^69^ mutants zebrafish were raised as incross and genotyped by PCR to identify homozygous *tp53^−/−^* or genotyped by Sanger sequencing to identify homozygous *atm^−/−^* mutants. *atm^−/−^; Tg (fabp10a:hUHRF1-EGFP ^mss1^* here after called *atm^−/−^; hUHRF1*, or *tp53^−/−^; Tg (fabp10a:hUHRF1-EGFP ^mss1^* here after called *tp53^−/−^; hUHRF1,* were generated by crossing *atm^−/−^; hUHRF1* or *tp53^−/−^; hUHRF1* adults to *atm^−/−^* or *tp53^−/−^*, respectively.

### EdU incorporation assay

hUHRF1 and control larvae at indicated time points were incubated in 250 μl of embryo medium containing 10% DMSO and 0.5 mM EdU (Thermo Fisher Scientific, resuspended in embryo water) for 20 minutes on ice. After 20 minutes, 50 ml of embryo medium was added to the larvae and then incubated at 28 °C for 30 minutes. For 7 and 10 dpf larvae, 0.22 μm filtered system water was used instead of embryo medium for the EdU incorporation. After EdU, larvae were fixed with 4% paraformaldehyde overnight at 4 °C, washed once with PBS and gradually dehydrated in 100% methanol and stored at 4 °C. EdU pulsed larvae were stained by click-IT (Thermo Fisher Scientific) as previously described^70^. Briefly, larvae were rehydrated in 100 % PBS and incubated twice with PBS containing 8 mM CuSO_4_ (Sigma Aldrich), 4 nM Sodium Azide (A555 or A647, Thermo Fisher Scientific) and 50 mM Ascorbic Acid (Sigma Aldrich) for 30 minutes at room temperature. Livers were dissected, stained with Hoechst (1:2000, Thermo Fisher Scientific) and GFP booster (1:1000, Chromotek) for 1 hour at room temperature before imaging. A full z-stack of each liver (z = 0.33μm) was acquired at Leica Stellaris 8 Confocal Microscope by using Lightening function. Images were quantified by Imaris (spot function) to count Hoechst, EGFP and EdU positive cells.

### Protein extraction and western blot

For protein extraction, 40-80 livers from hUHRF1 and controls were manually dissected and collected in ice-cold PBS. After centrifugation, livers were resuspended in RIPA buffer (Sigma Aldrich, 2 μl of RIPA buffer each liver) supplemented with protease inhibitors cocktail (Roche), incubated 15 minutes of ice and sonicated to release proteins with a Hand sonicator (Hielscher Ultrasonics GmbH). Protein lysates were cleared by centrifugation, supplemented with Leammli buffer (BioRad), boiled for 5 minutes at 95 °C, loaded into 10% acrylammide/bisacrylammyde denaturing gel (Sigma Aldrich) and run at 80 V. Proteins were transferred onto PDVF membrane (Thermo Scientific) for 1.5 hours at 300 mA on ice. Membrane was blocked for 1 hour at room temperature with TBS (BioRad) supplemented with 0.1% Tween-20 (Sigma Aldrich) containing 5% skim milk (Sigma Aldrich), incubated overnight at 4 °C with 1: 1000 primary antibody diluted in blocking solution (H3K9me3, Active Motif; H3, Santa Cruz) followed by secondary antibody diluted 1: 2500 in blocking solution (HRP-conjugated anti-Rabbit, Promega). Chemiluminescence was revealed with ECL Clarity (BioRad) and imaged with ChemiDoc MP Imaging System (BioRad). Images were quantified with GelAnalyzer.

### Immunofluorescence

hUHRF1 and control larvae from indicated time points were collected and fixed with 4 % PFA at 4 °C overnight. After fixation, larvae were washed twice with PBS. Livers were manually dissected and incubated with PBS containing 1 % Triton-X (Sigma Aldrich) and 20 μg/ml of Proteinase K (Machery-Nagel) for 10 minutes at room temperature to improve permeabilization. After permeabilization, livers were washed 3 times with PBS and incubated 1 hour at room temperature with blocking solution (PBS containing 2 % BSA, Sigma Aldrich) followed by incubation at 4 °C overnight with primary antibody staining (blocking solution containing 1:100 primary antibody). Primary antibody used: H3K9me3 (#39161, Active Motif), yH2AX (gtx127342, Genetex), pRPA (ab211877, AbCam). After primary antibody staining, livers were washed 3 times with PBS containing 0.1% Tween-20 (Sigma Aldrich) followed by 2 washed with PBS, and secondary antibody staining (blocking solution with 1:300 secondary antibody and 1:2000 Hoechst, Thermo Fisher Scientific). Secondary antibody used: anti-rabbit A 555 (Thermo Fisher Scientific). After secondary antibody staining, livers were washed 3 times with PBS, mounted with vectashield (Vector), imaged at Leica Stellaris 8 Confocal Microscope and quantified with LasX.

### RNA extraction, RNAseq and bulk RNAseq analysis

For RNA extraction, 20-30 livers were manually dissected from 5 independent clutches of hUHRF1, p53^-/-^, p53^-/-^;hUHRF1 and controls and collected directly in 500 μl of Trizol (Thermo Fisher Scientific). RNA was extracted by following manufacturer’s instruction with some modification. After chloroform, RNA was precipitated in isopropanol (Sigma Aldrich) with 10 μg of Glycoblue (Thermo Fisher Scientific) overnight at – 20 °C. After precipitation, RNA was centrifuged at 4 °C for 1 hour and resuspended in 20 μl of DNAse/RNAse free water. Possible genomic DNA contamination was removed with RapidOut DNA removal Kit (Thermo Fisher Scientific) for 30 minutes at 37 °C. 100 ng of RNA was used for library preparation with ILMN Strnd Total RNA Lig w/RBZ+ (Illumina). Libraries were sequenced on NovaSeq Illumina platform. Quality of the sequences was assessed by using MultiQC v1.0 (https://multiqc.info). After adaptor removal and trimming, reads were aligned to the D. rerio GRCz10 reference genome, with manual insertion of UHRF1 and EGFP sequence, using HISAT2 with default parameters^71^, mapped and counted with HTSeq^72^. Differential gene expression was calculated using a generalized linear model implemented in DESeq2 in Bioconductor^73^ to test differential gene expression between hUHRF1 transgenic livers and WT sibling controls. Adjusted p-value with a false discovery rate of <0.05 was used to determine significantly differentially expressed genes between transgenics and controls.

Quantification of repetitive elements (REs) were assessed by SQuIRE workflow^74^ to quantify expression at the subfamily level and properly assign multi-mappers sequences (due to the nature of repetitive elements) using expectation-maximization (EM) algorithm. In brief, trimmed fastq were aligned to D. rerio GRCz10 reference genome using STAR. REs annotation from RepeatMasker was used for alignment with unique mapping reads aligned to a single locus and marking multi-mapped reads to multiple locations and further refined with EM algorithm with “auto” parameter. Total counts (unique and multi-mapped) are aggregated by subfamily of REs and the output count table from SQuIRE has been analyzed by DEseq2 with standard parameters to calculate differential expression of REs between condition.

### DNA extraction, RRBS and DNA methylation analysis

Genomic DNA was extracted from 20 to 100 livers from 3 independent clutches by using a DNA extraction buffer (10 mM Tris-HCl pH9, 10 mM EDTA, 200 mM NaCl, 0.5% DSD, 200 μg/ml proteinase K) as previously described^65^. DNA was resuspended in water and quantified by Qubit dsDNA High Sensitivity kit.

RRBS was performed on ∼80 ng of genomic and digested with 200 U of MspI (New England Biolabs) and used for preparing library, as previously described^75^, with some modifications. To avoid loss of gDNA, after MspI digestion, end repair, and A-tailing, the reactions were stopped by heat inactivation. The adaptors used for multiplexing were purchased separately (Next Multiplex Methylated Adaptors-New England Biolabs). Libraries were size-selected by dual-step purification with Ampure XP Magnetic Beads (Beckman Coulter) to specifically select a region of fragments from 175 bp to 670 bp. Bisulfite conversion was performed with Lightning Methylation Kit (ZYMO Research) by following the manufacturer’s instructions. Libraries were amplified using KAPA HiFi HotStart Uracil+ Taq polymerase (Roche) and purified with Ampure XP Magnetic Beads (Beckman Coulter) before sequencing. Libraries were sequenced using the Illumina Nextseq 555 platform. Quality control of the RRBS sequencing data was assessed using FASTQC and Trimmomatic and aligned to the reference genome GRCz10 as described previously^76^.

RRBS data was analyzed for CpG methylation levels using the R package ‘methylKit’^77^. CpGs covered at least 10 times in one biological replicate were included in the analysis. CpGs with a q-value with a false discovery rate of <0.01 were considered differentially methylated. Genomic element annotation of CpGs was performed with R package ‘genomation’. For plotting and statistical analysis, R package ‘ggplot’ and GraphPad Prism software were used.

### Preparation of single cell suspension and scRNAseq

150-800 livers for each time point from Controls and hUHRF1 were manually dissected and put in ice-cold PBS containing 2 % FBS (Gibco) and 1 % penicillin/streptomycin (Gibco). After dissection, livers were briefly centrifuged, resuspended in 1 ml of Trypsin/EDTA 0.05% (Sigma Aldrich) and incubated at 28 °C for 12-20 minutes. Every 2 minutes livers were pipetted 10 times to facilitate cell dissociation until all the cells resulted in a homogenous suspension. 200 μl of FBS (Gibco) was added to inactivate Trypsin and cells were centrifugated at 4 °C for 7 minutes at 500 g. After centrifugation cells were washed 3 times with PBS containing 0.4 % BSA (Sigma Aldrich). Cells were resuspended in 50-100 μl of PBS containing 0.4% BSA and counted with a Burker Chamber. Cell suspension at a concentration of 600-700 cells/μl was used to load Chip for 10X Genomics acquisition. All steps of single-cell sequencing were performed following the Chromium Next GEM Single Cell 3ʹ Reagent Kits v3.1 (Dual Index).

### Single-Cell Data analysis

Single-cell RNA sequencing (scRNA-Seq) data were processed and filtered using single-cell analysis package Seurat v5.0.1. Initial quality control and filtering of scRNA-Seq data were performed to ensure the retention of high-quality cells. In that line, cells with fewer than 200 detected features and nCount_RNA or mitochondrial gene content greater than 20% were excluded from downstream analysis. Due to the large number of cells in the 10 dpf CAAX sample in comparison to other samples, we downsampled the dataset before data filtering to include only 10,000 randomly selected cells, with a fixed random seed (123) to ensure reproducibility. Data filtering resulted in 12643 cells for 5 dpf Controls, 7684 cells for 7 dpf Controls, 8206 cells for 10 dpf Controls, 6841 cells for 14 dpf Controls, 10409 cells for 5 dpf hUHRF1, 5265 cells for 7 dpf hUHRF1, 9631 cells for 10 dpf hUHRF1, 8144 cells for 14 dpf hUHRF1 and 8260 cells for 20 dpf hUHRF1 corresponding to a total of 77083 CAAX and hURFF1 cells to be integrated. Prior to data integration, each sample was independently normalized and scaled with a with scaling factor of 10,000. Highly variable features were identified using the “vst” method with the top 2,000 features selected. Data integration was performed with Seurat package using the FindIntegrationAnchors function with canonical correlation analysis (CCA)^78^. Batch effects were corrected based on sample identity. Dimensionality reduction and clustering of scRNA-Seq data were performed using principal component analysis (PCA), retaining the top 25 principal components. Uniform Manifold Approximation and Projection (UMAP) dimensionality reduction was applied using these components for visualization. Neighbor identification was conducted with FindNeighbors (k = 20), followed by clustering using FindClusters with a resolution of 0.4. Differentially expressed genes (DEGs) in each cell cluster were identified using FindMarkers function, and statistically significant markers between compared groups were selected based on an adjusted p-value threshold (p-value < 0.05). All dimensionality reduction and DEG analyses were conducted using the Seurat package.

### Analysis of publicly available human scRNA-Seq datasets

Single-cell RNA sequencing data from 10 hepatocellular carcinoma (HCC) patients were obtained from the publicly available dataset GSE149614^50^. Raw gene expression counts were analyzed and preprocessed using the Seurat package (version 5.0.1), which included normalization, identification of highly variable genes, and scaling of the data. Dimensionality reduction was performed using PCA, followed by t-SNE for visualization of cell populations. Cell clustering was performed using the FindClusters function with a resolution of 3.

### Pseudotemporal Dynamics of Tumor Evolution

To identify trajectories that could represent cellular pseudotemporal paths resulting in HCC development we utilized the Monocle3 R package (version 1.3.7). Initially, the Seurat object was transformed into a CellDataSet (cds) object with the SeuratWrappers package (version 0.3.5). Subsequently, the learn graph function from Monocle3 was applied to infer the global trajectory structure by applying reversed graph embedding algorithms. Pseudotime ordering of cells was then performed using the order_cells function, whereas Progenitor2 cells were designated as the root nodes of the trajectory. Differentially progressing genes along the pseudotime trajectory were identified using the graph_test function. To cluster the 100 differentially progressing genes based on their pseudotemporal expression levels, a smoothing spline with three degrees of freedom was fitted to the expression data for each gene. Finally, z-score normalization was applied to standardize expression values, enabling consistent comparison of expression patterns across genes.

### Lineage tracing

fabp10a:BB-NTR and fabp10a:CreER line were crossed to generate fabp10a:BB-NTR; fabp10a:CreER and crossed to hUHRF1; fabp10a:CreER. Embryos at 3 dpf were screened for the presence of the cassettes by using the transgenesis markers (“red eye” for fabp10a:BB-NTR and “green eye” for fabp10a:CreER) alone or together with hUHRF1. hUHRF1; fabp10a:BB-NTR; fabp10a:CreER and fabp10a:BB-NTR; fabp10a:CreER were treated at 80 hpf with 10 μg/ml of 4- Hydroxytamoxifen (MedChem Express) for 24 hours. After treatment, larvae were washed and raised till 20 dpf with regular feeding. At 6 and 20 dpf larvae were fixed in 4 % PFA at 4 °C overnight, transferred in PBS. Livers were manually dissected and mounted on glass slides with Vectashiled (Vector). Images were acquired at Leica Stellaris 8 Confocal Microscope.

### Analysis of cell heterogeneity

hUHRF1 and nls-mCherry lines were crossed and hUHRF1;nls-mCherry or nls-mCherry larvae were collected at indicated time points and fixed in 4 % PFA at 4 °C overnight and then transferred in PBS. Livers were manually dissected, permeabilized with PBS containing 1 % Triton-X (Sigma Aldrich) and 20 μg/ul Proteinase K (Machery-Nagel) for 10 minutes at room temperature and stained with 1:2000 Hoechst (Thermo Fisher Scientific) in PBS for 1 hour at room temperature. Livers were mounted on a glass slide with Vectashield (Vector) and images were acquired with Leica Stellaris 8 Confocal Microscope. For each liver, the lasers were set to have 1-2 saturated cells for each channel. Image analysis is performed with Fiji by creating a mask on Hoechst channel and quantify the intensity of mCherry and EGFP for all the cells present in the mask. For each image, for each channel, levels of intensity of each cell were normalized based on the cell with highest intensity in that image and relative intensity was plotted for visualization in Graph Prism 8.

## Statistical analysis

Statistical analysis was performed in GraphPad Prism 8. Number of replicates for each experiment are indicated in each figure legends. Methods to evaluate the statistical significance include two-tailed Student’s *t*-test with adjustment for multiple comparisons, t-test, or Chi-square for categorical variables. Tests used are indicated in figure legend. All the plots were generated in GraphPad Prism 8 and RStudio 3.3.1.

