## Supplementary material for "Epigenetic Disordering Drives Stemness, Senescence Escape and Tumor Heterogeneity": Supplemental Material.pdf

### Supplemental Material and Methods

#### Crispr/Cas9 deletion of human UHRF1

sgRNA targeting the human UHRF1 gene was produced by sgRNA IVT kit (Takara Bio), the resulting RNA was isolated using Trizol (Invitrogen) and was quantified by Qubit. The sgRNA was diluted to 50 ng/μL, mixed with an equal volume of previously diluted nls-Cas9 protein (IDT; 0.5 μl of nl-Cas9 added with 9.5 μL of 20 mM HEPES; 150 mM KCl, pH 7.5) and incubated at 37 °C for 5 min. We then injected 1 nl into 1–2 cell stage embryos from an incross of hUHRF1 which were then raised to 120 hpf. At 120 hpf embryos were screened for the presence of EGFP in the liver in both not injected and injected Controls. 75 % positive is the expected percentage of EGFP in not-injected controls while the injected, if crispants, should show a decreased percentage. EGFP positive and negative from each group were used to perform SA-ß-galactosidase staining.

#### Mapping fabp10a:hUHRF1-EGFP integration

gDNA from hUHRF1-EGFP positive zebrafish was extracted as described. 5 μg of gDNA was digested with AluI (New England Biolabs) at 37 °C for 3 hours. Digested DNA was purified with PCR purification kit (Sigma Aldrich) then ligated with T4 DNA ligase (New England Biolabs) at 16 °C overnight. Ligated products were PCR amplified with PCR followed by nested PCR by targeting the 5' and 3' region of the cassette directed outside the cassette. Water was used as PCR control and WT gDNA was processed in parallel as negative control. PCR product was run on a 1 % agarose gel and interested band was gel purified with Gel Elute kit (Sigma Aldrich). Purified products were sequenced and mapped to zebrafish genome by nBLAST.

#### In situ hybridization

6 dpf hUHRF1 and wild type larvae were fixed with 4 % PFA at 4 °C overnight. After fixation larvae were gradually de-hydrated into 100 % methanol and incubated overnight for at 4 °C. Larvae were gradually re-hydrated and in situ hybridization was performed as per manufacturer's instruction (Molecular Instruments).

### Supplemental Figure Legends

**Supplemental Figure 1. hUHRF1 phenotype does not depends on integration site. A.** Schematic representation of human UHRF1 functional domains. Arrows indicates the location of the sgRNA sequence. **B.** Stack bar of percentage of EGFP positive larvae (indicating functional hUHRF1) in not injected controls and embryos injected with sgRNA 1 or 2. **C.** Stack bar of percentage of SA-ß-galactosidase positive larvae not injected controls and embryos injected with sgRNA 1 or 2, divided by EGFP presence. **D.** Location of integration of Tg(fabp10a:hUHRF1-EGFP) cassette. **E.** Histogram of normalized counts of genes surrounding the Tg(fabp10a:hUHRF1-EGFP) cassette at 80 hpf and 120 hpf.

**Supplemental Figure 2. RNAseq of 80 hpf hUHRF1 compared Controls. A.** MA plot of log2 fold change calculated on hUHRF1 compared to Controls livers at 80 hpf. **B**. Volcano plot of Log2 fold change calculated on hUHRF1 compared to Controls livers at 80 hpf. **C.** Venn diagram of differentially expressed genes (padj < 0.05) in hUHRF1 compared Controls at 80 hpf and 120 hpf.

**Supplemental Figure 3. Validation of Atr inhibition. A.** Treatment scheme of Atr inhibitor as performed in hUHRF1 and Control larvae. To determine the efficacy, H2O2 was used as DNA damage inducer. **B.** Western blot of Control and H2O2 treated larvae in presence and absence of the Atr inhibitor VE-821.

**Supplemental Figure 4. RNAseq of 120 hpf hUHRF1 compared Controls and tp53-/-; hUHRF1 compared to tp53-/-. A.** MA plot of log2 fold change calculated on hUHRF1 compared to Controls livers at 120 hpf. **B**. Volcano plot of Log2 fold change calculated on hUHRF1 compared to Controls livers at 120 hpf. **C.** MA plot of log2 fold change calculated on hUHRF1 compared to Controls livers at 120 hpf. **D**. Volcano plot of Log2 fold change calculated on tp53-/-;hUHRF1 compared to tp53-/- livers at 120 hpf. **E.** Venn diagram of differentially expressed genes (padj < 0.05) in hUHRF1 compared Controls and tp53-/-;hUHRF1 compared to tp53-/-.

**Supplemental Figure 5. scRNAseq of hUHRF1 and Controls at 5 dpf. A.** Heatmap of top 10 differentially expressed genes in each population identified at 5 dpf compared to genes present in the whole dataset. **B.** Violin plot of EGFP expression levels across population in Controls. **C.** Violin plot of EGFP expression levels across population in hUHRF1.

**Supplemental Figure 6. EMT markers in scRNAseq.** Box plot of expression EMT markers in Mesenchymal cells (Mesenchymal 1 and 2) in Controls and hUHRF1 at 5 dpf.

**Supplemental Figure 7. hUHRF1 heterogeneity in hUHRF1 livers. A.** Representative images of Hoecsht, EGFP and nls-mCherry in hUHRF1;mCherry and mCherry fish at 5 dpf. **B.** Quantification of Relative Intensity calculated on maximum intensity level of each image in mCherry in nls-mCherry livers at different time points. **C.** In Situ Hybridization of human UHRF1 mRNA in 6 dpf hUHRF1 livers.

**Supplemental Figure 8. Cell identity genes are enriched in distinct populations of cells identified by scRNAseq of hUHRF1-high and Controls livers. A.** Heatmap of top 10 DEGs in each population identified at 5, 7, 10, 14, 20 dpf compared to genes in the whole dataset. **B.** Stack bar showing the number of cells present in hUHRF1 and controls for each population at different time points. Note that there was no sample sequenced for controls at 20 dpf and therefore the comparison with controls is missing for this time point.

**Supplemental Figure 9. EGFP across population in scRNAseq. A.** EGFP expression levels in different hepatocyte populations across time points. **B.** Gaussian distribution of EGFP expression levels in hepatocytes population, biliary epithelial cells, progenitor and mesenchymal cells in Controls and hUHRF1 and cut-off used to divided cells in top, middle, and low.

**Supplemental Figure S10. Stem cell genes are upregulated in progenitor cell populations in hUHRF1-high livers during hepatocarcinogeneis.** Feature plot of *epcam*, *tgfb1a*, *klf4* and *prom1a* in all cell population across time points in controls and hUHRF1-high samples.

**Supplemental Figure S11. Tamoxifen effectively eliminates BFP expression in hepatocytes at 6 dpf.** Representative images of 6 dpf livers of BB-NTR Controls untreated (**A**), BB-NTR controls treated with Tamoxifen at 3.5 dpf for 24 hpf (**B**) and hUHRF1; BB-NTR treated with Tamoxifen at 3.5 dpf for 24 hpf (**C**).

**Supplemental Tables**

**Supplemental table S1. Bulk RNAseq at 80 hpf.** Significantly (-adj < 0.05) differentially expressed genes (-adj < 0.05) in bulk RNAseq at 80 hpf in hUHRF1 compared wild type controls.
**Supplemental table S2. Bulk RNAseq at 120 hpf.** Significantly (-adj < 0.05) differentially expressed genes (-adj < 0.05) in bulk RNAseq at 120 hpf in hUHRF1 compared wild type controls.

**Supplemental table S3. Bulk RNAseq at 120 hpf in tp53-/-.** Significantly (-adj < 0.05) differentially expressed genes (-adj < 0.05) in bulk RNAseq at 120 hpf in tp53-/-;hUHRF1 compared to tp53-/-.

**Supplemental table S4. scRNAseq 5 dpf.** Differentially expressed genes of scRNAseq of each population compared all cells at 5 dpf.

**Supplemental table S5. scRNAseq 5, 7, 10, 14, 20 dpf.** Differentially expressed genes of scRNAseq of each population compared all cells at all time points.
