## Supplementary figures and images for "Epigenetic Disordering Drives Stemness, Senescence Escape and Tumor Heterogeneity"

### 07_Supplemental Figure S1.png

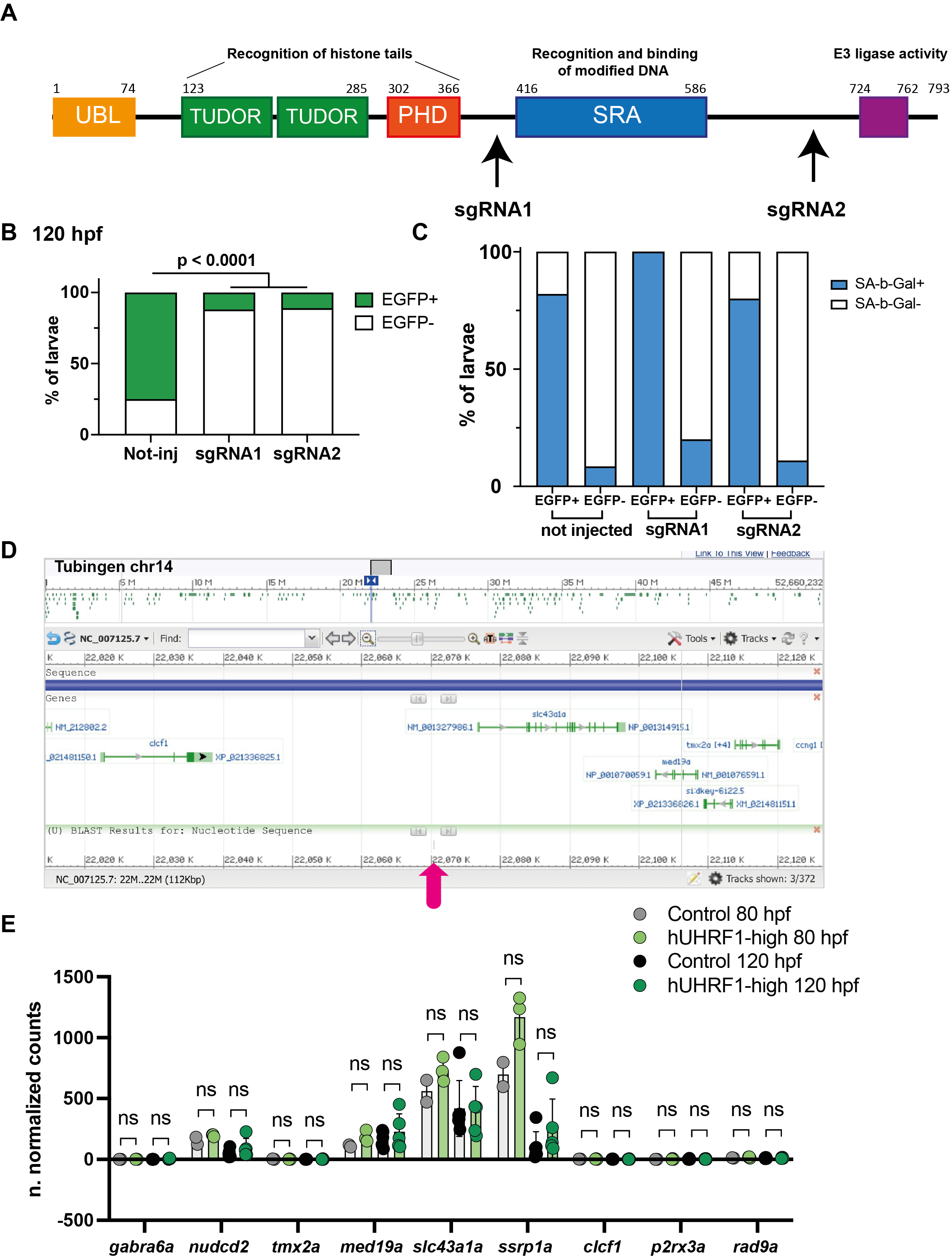

### 08_Supplemental Figure S2.png

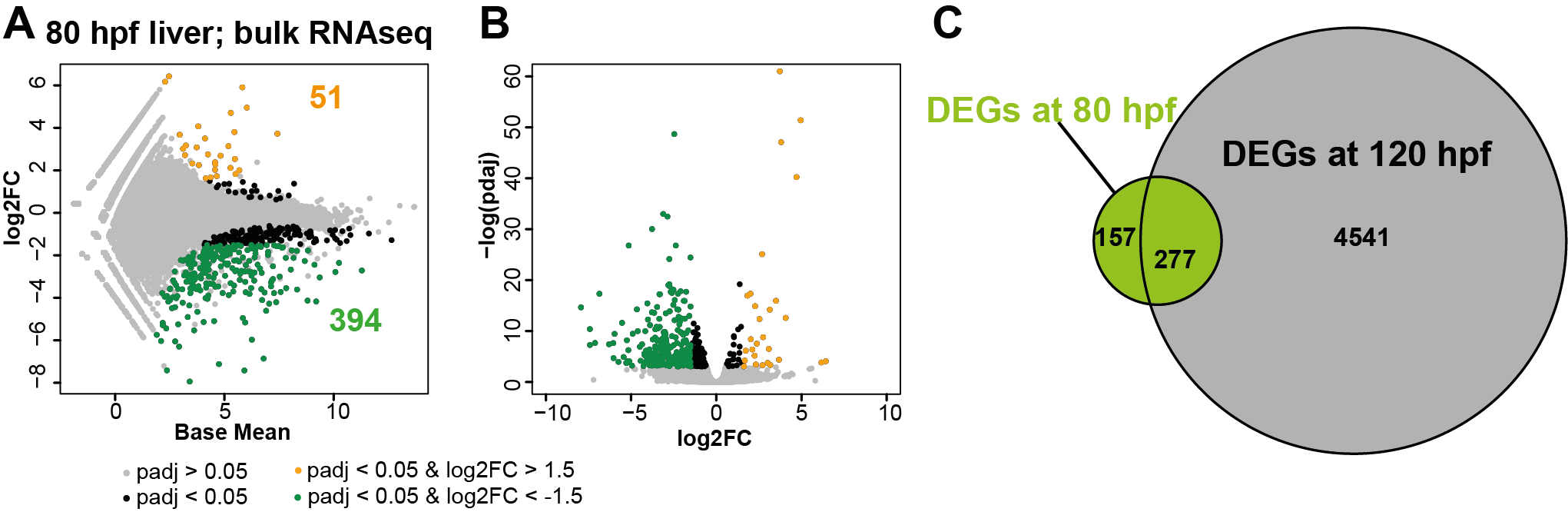

### 09_Supplemental FIgure S3.png

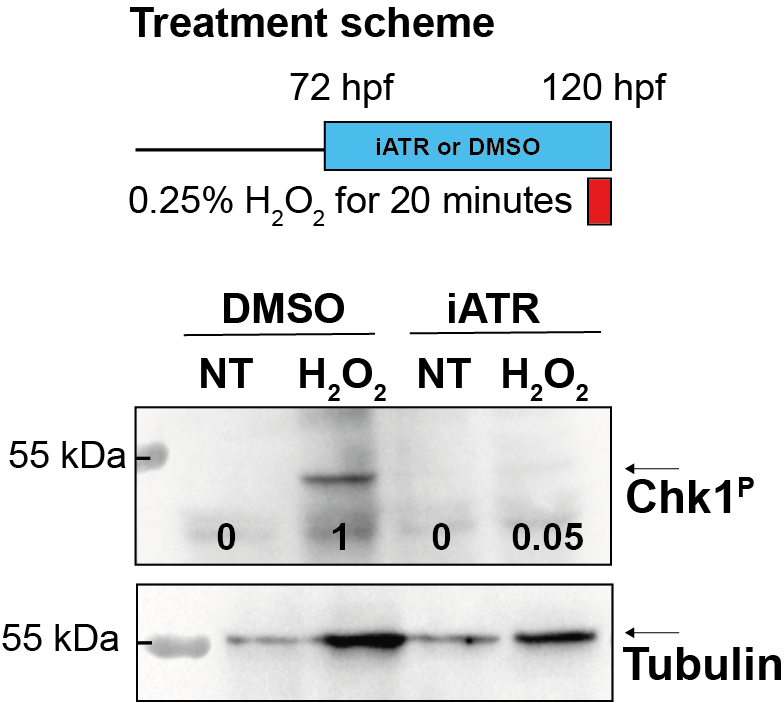

### 10_Supplemental Figure S4.png

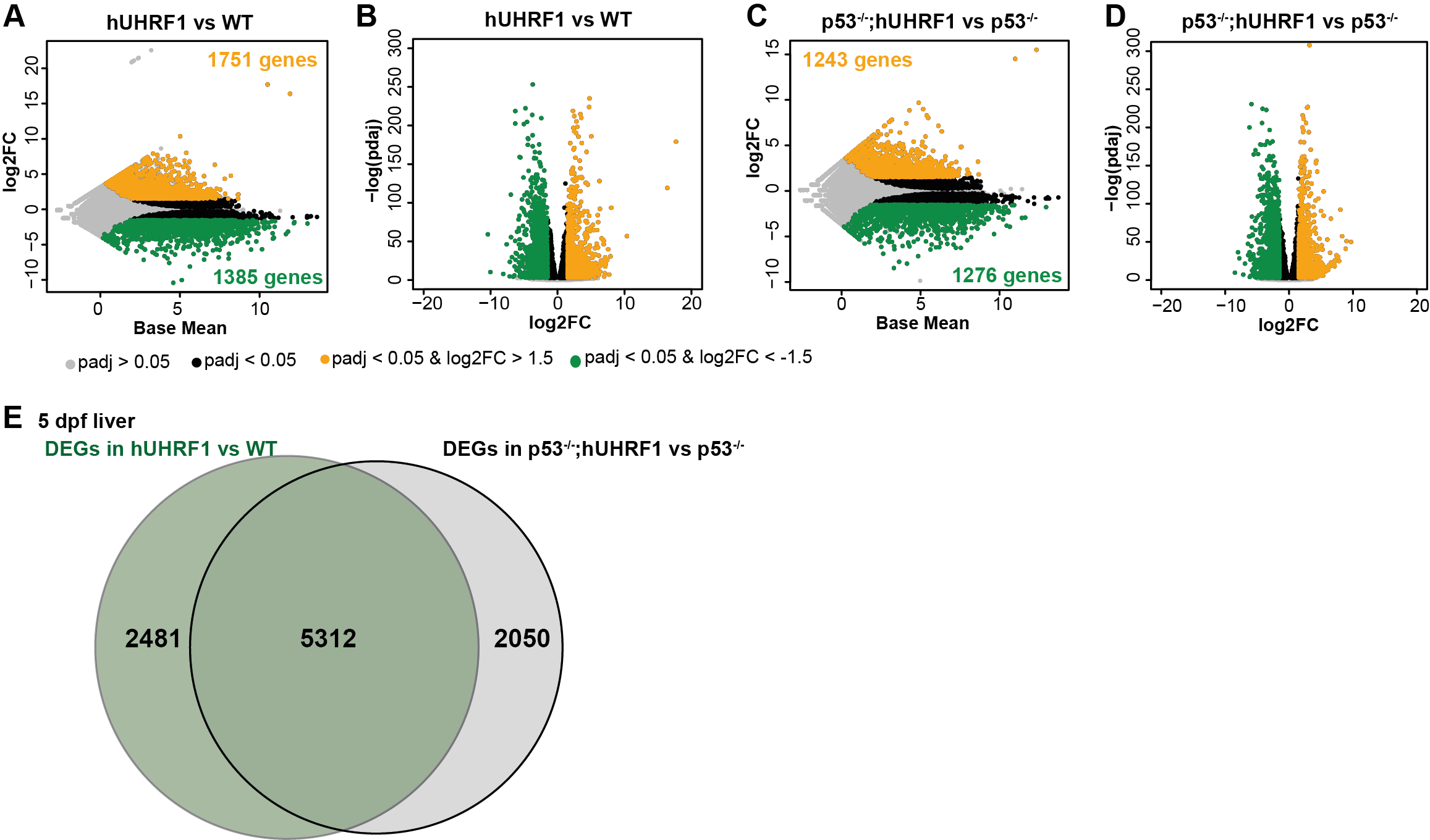

### 11_Supplemental Figure S5.png

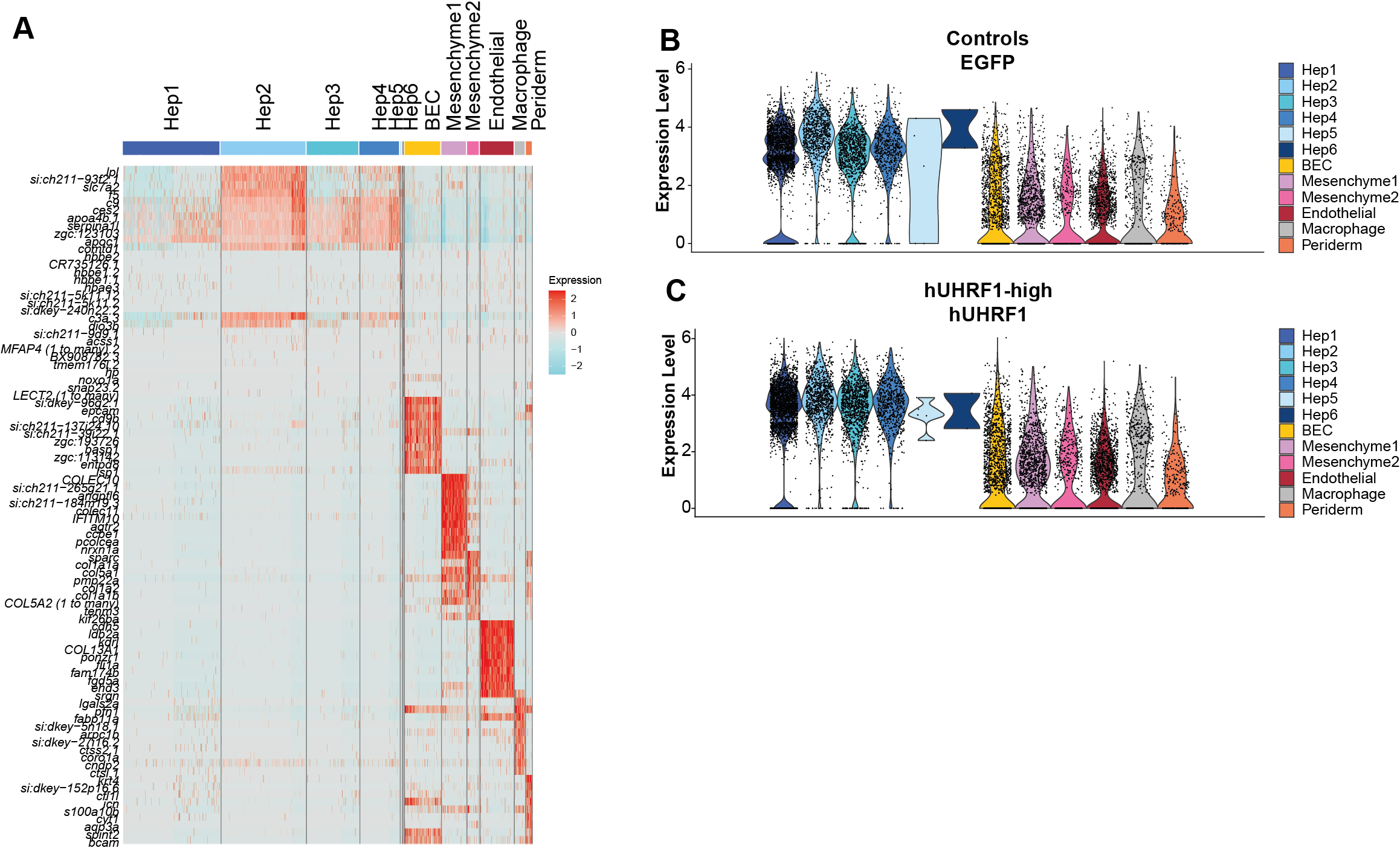

### 12_Supplemental Figure S6.png

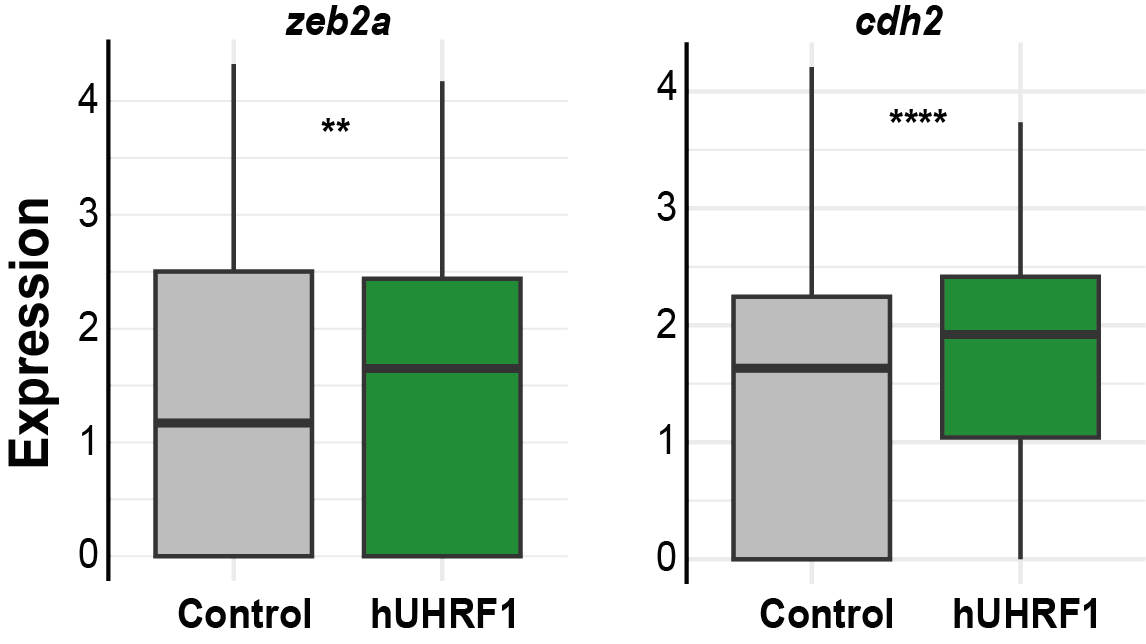

### 13_Supplemental Figure S7.png

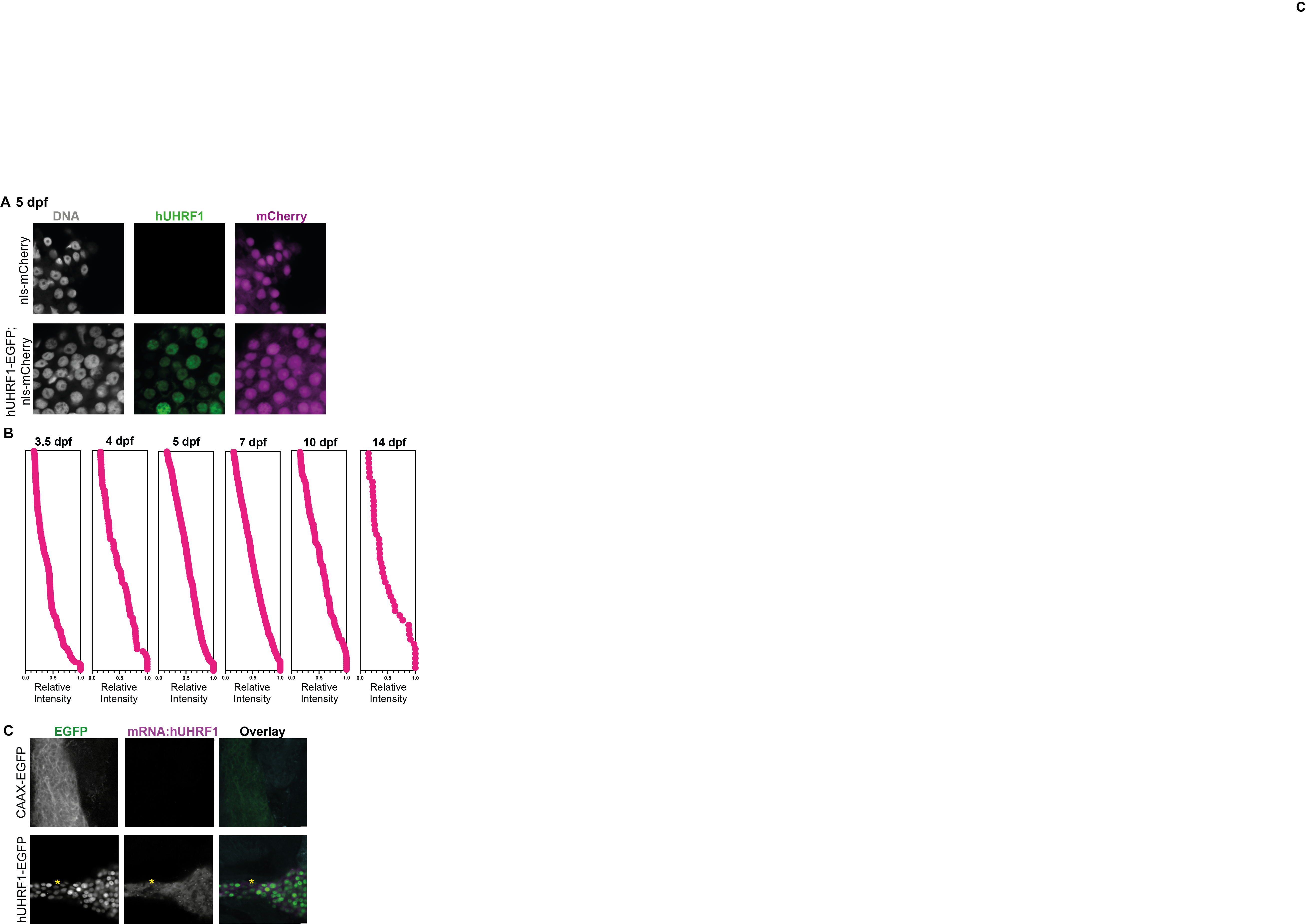

### 14_Supplemental Figure S8.png

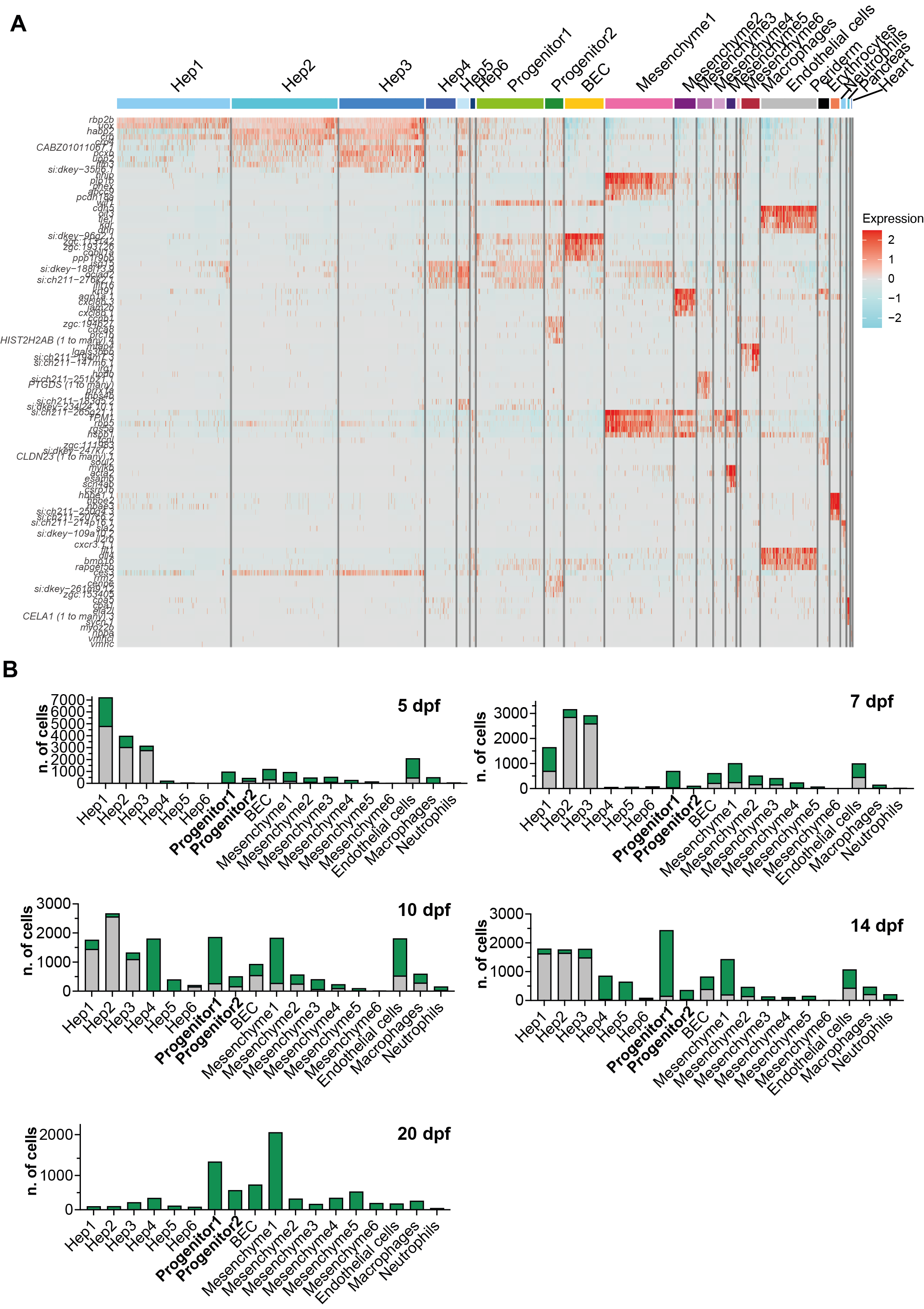

### 15_Supplemental Figure S9.png

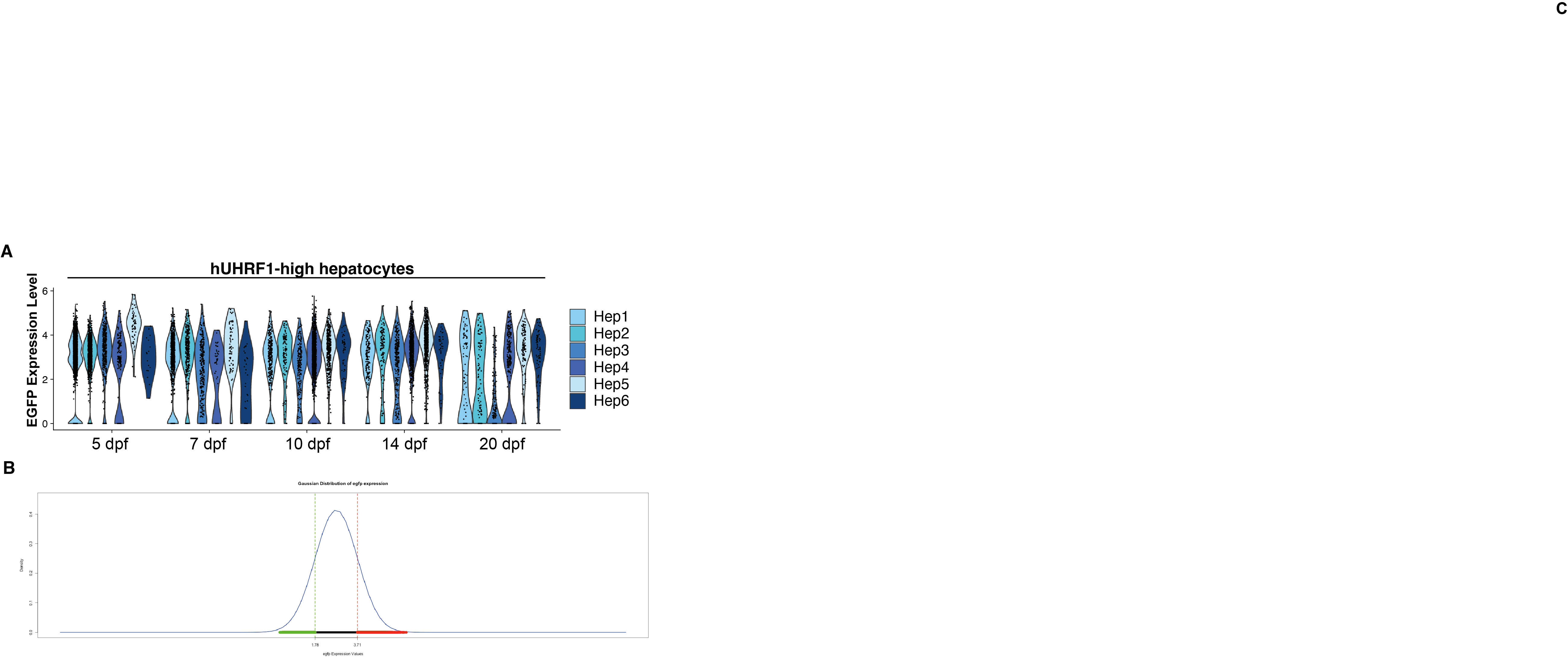

### 16_Supplemental Figure S10.png

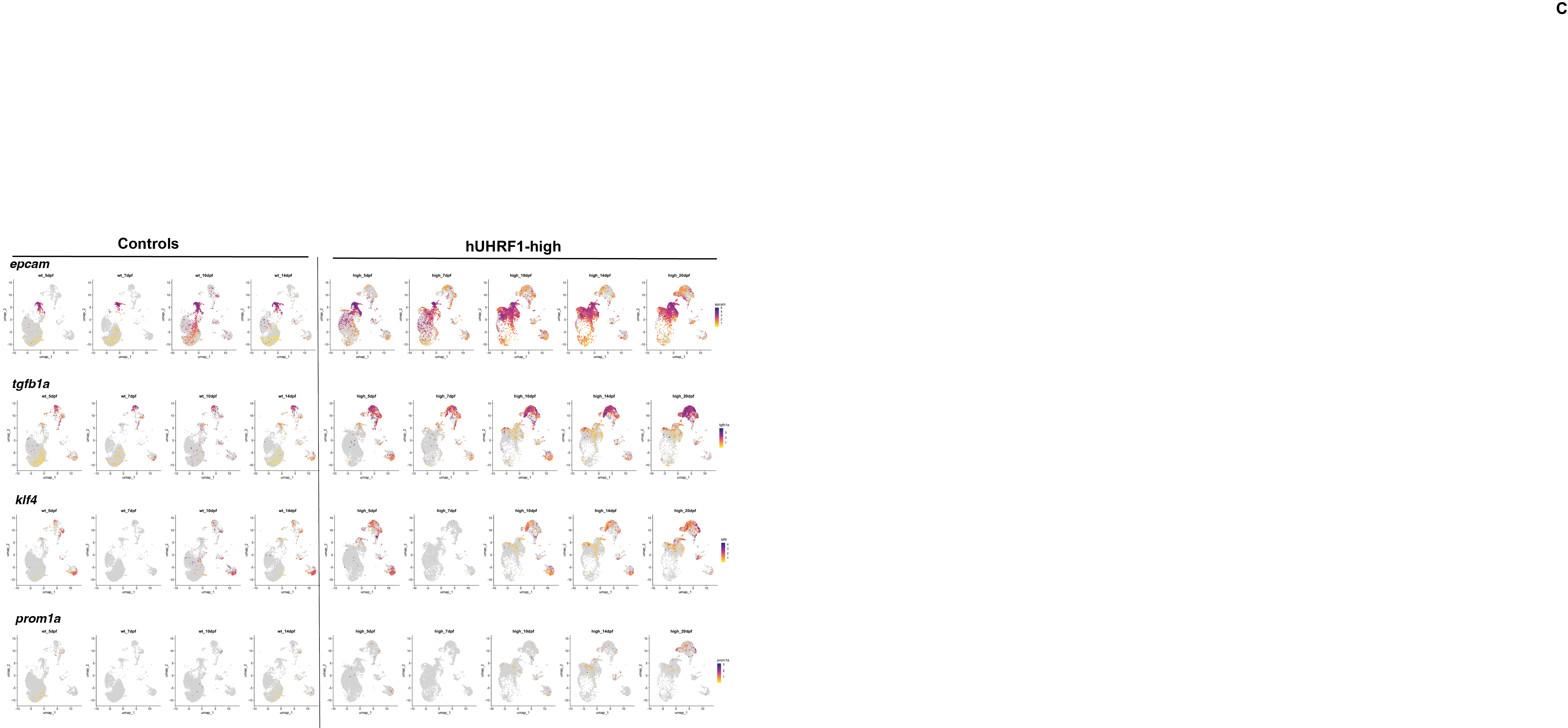

### 17_Supplemental Figure S11.png

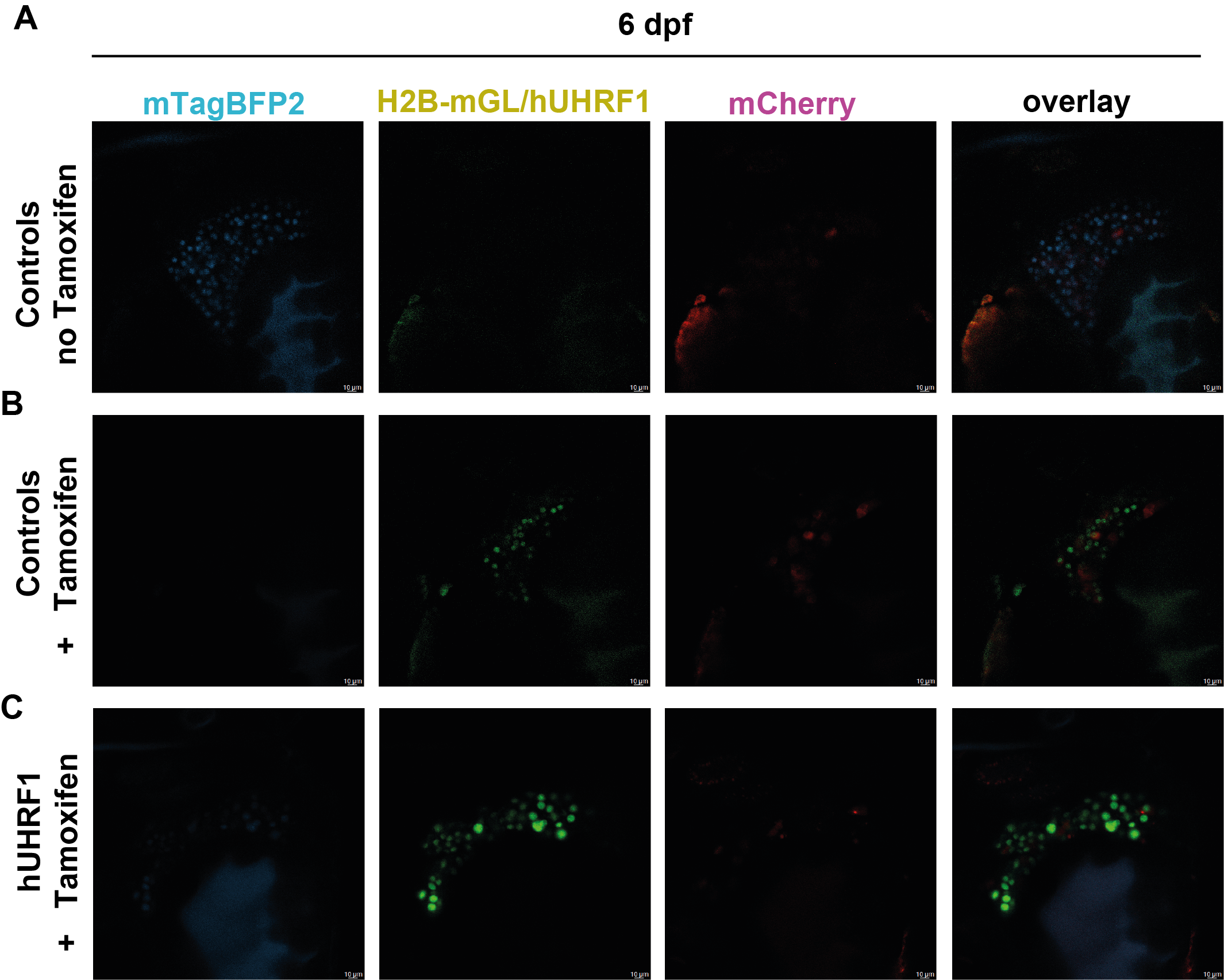
